# Confinement modulates axial patterning in regenerating Hydra

**DOI:** 10.1101/2024.06.13.598813

**Authors:** Yonit Maroudas-Sacks, Liora Garion, S Suganthan, Marko Popović, Kinneret Keren

**Affiliations:** Department of Physics, Technion– Israel Institute of Technology, Haifa 32000, Israel; Max-Planck Institute for Physics of Complex Systems, MPI-PKS, Nothnitzer Str. 38, Dresden, 01187, Germany; Cluster of Excellence, Physics of Life, Technische Universitat Dresden, Arnoldstrasse 18, Dresden, 01307, Germany; Center for Systems Biology, Pfotenhauerstrasse 108, Dresden, 01307, Germany; Network Biology Research Laboratories and Russell Berrie Nanotechnology Institute, Technion– Israel Institute of Technology, Haifa 32000, Israel

## Abstract

The establishment of the body plan is a major step in animal morphogenesis. The role of mechanical forces and feedback in patterning the body plan remains unclear. Here we explore this question, by studying regenerating Hydra tissues confined in narrow cylindrical channels which constrain their morphology. We find that frustration between the orientation of the channel and the inherited axis in the regenerating tissues can lead to the formation of a multiaxial body plan. The morphological outcome is directly related to the pattern of nematic topological defects that emerges in the organization of the supracellular actomyosin fibers. When the inherited axis, which can be read out from the initial alignment of the supracellular fibers in the confined spheroid, is parallel to the channel’s axis, the tissue regenerates normally into animals with a single body axis aligned with the channel. However, regenerating spheroids that are confined in a frustrated perpendicular configuration often develop excess defects (including negatively-charged -½ defects) and regenerate into multiaxial morphologies. The influence of mechanical constraints on the regenerated body plan argues against an axial patterning mechanism that is based solely on inherited gradients of biochemical morphogens. We further show that the dependence of the regeneration outcomes on the initial tissue orientation can be recapitulated by a biophysical model which considers the coupled dynamics of the nematic organization of the actomyosin fibers and a morphogen concentration field, incorporating a mechanochemical feedback loop involving strain-dependent morphogen production at defect sites.

## INTRODUCTION

The emergence of the body plan during morphogenesis is a prime example of biological pattern formation that is still poorly understood. Research on morphogenesis usually emphasizes the role of gradients of biochemical signaling molecules (morphogens), together with positional-sensitive response of cells, in defining the body axes that emerge during development [1]. However, a growing number of studies, show that mechanical forces and feedback also play an important role in patterning the body plan in developing animals [2-4]. These findings suggest that patterning during morphogenesis arises through a self-organization process that couples the dynamics of biochemical morphogens with mechanical fields through various mechanochemical feedback loops. Despite much effort, understanding the coupling between the biochemical signaling pathways and tissue mechanics in animal morphogenesis remains an outstanding challenge. Here we investigate this question by applying an external constraint on regenerating Hydra, and examining how this mechanical perturbation influences the patterning of the body plan.

Hydra is a small fresh-water predatory animal that has a simple uniaxial body plan and exhibits remarkable regeneration capabilities [5]. Bisected Hydra regenerate a head or foot according to their original polarity [6], and even small excised tissue pieces [7] or aggregates of dissociated cells [8] regenerate into complete animals within a couple of days. Hydra has been a classic model system for studying axial patterning in animal morphogenesis for over a century, from the pioneering work of Ethel Brown who discovered the head organizer in Hydra [9, 10]. Subsequent work established the presence of graded activation and inhibition properties in the tissue [11, 12] and uncovered many of the molecular players involved in setting up the body plan [13-16]. However, our understanding of the mechanisms underlying axial patterning in Hydra remains limited [17].

Our previous work focused on the role of the supracellular arrays of actomyosin fibers in guiding and stabilizing the body axis in regenerating Hydra [18-21]. Mature Hydra present a nematic array of parallel contractile actomyosin fibers called myonemes with a characteristic pattern of topological defects in the fiber alignment field [20]. In the ectoderm, where the fibers are aligned along the body axis, we find aster-shaped +1 defects at the tip of the head and foot regions. In addition, +1 nematic defects at the tip of each tentacle are balanced by two -½ nematic defects at the tentacle’s base. The endoderm fibers exhibit a similar pattern of defects, except that the fibers are aligned in a circumferential manner, orthogonal to the ectodermal fibers, so the +1 defects are vortex-shaped rather than asters. These fiber arrays function as muscles, facilitating the contraction and extension of the Hydra body as it moves around and preys [22].

During regeneration from excised tissue segments, we have shown that the alignment of the supracellular actomyosin fibers in the excised tissue confers a structural memory that persists during regeneration, and guides the formation of the body axis in the regenerated animal [19-21]. The excised tissue first fold into hollow spheroids that largely maintain their aligned fiber array, but undergo extensive reorganization near the closure region and lose the fiber organization there. Following the induction of nematic order across the regenerating tissue, defects in the nematic alignment of the fibers must emerge due to topological constraints [20]. We have shown that topological defects emerge in the initially disordered regions of the folded spheroid [20, 23]. Notably, the locations of defect formation can be identified from the actomyosin fiber pattern already at the onset of the regeneration process, and eventually become the sites of head and foot formation in the regenerated animal [23]. As such, our work suggests that the nematic actomyosin fiber orientation field with its characteristic set of topological defects is a relevant field for Hydra morphogenesis. How this mechanical field is coupled to the establishment of the body plan during Hydra morphogenesis is still unknown.

Application of external constraints on developing tissues can be informative for deciphering the role of mechanical constraints in normal development, but also more generally, for uncovering potential feedback mechanisms by introducing mechanical perturbations and studying how the system responds. Recent work has begun to explore the role of mechanical constraints in morphogenesis, both in developing animal in vivo and more recently in synthetic systems [4, 24, 25]. For example, *in vivo* mechanical perturbations revealed that oriented junction growth during *Drosophila* germ band extension is due to local polarized stresses driven by actomyosin contractions [26]. *In vitro*, confinement of cultured human embryonic stem cells to micro-patterned adhesive regions was sufficient to drive a self-organized pattern of cell differentiation into the 3 germ layers that arise during gastrulation [27]. More recently, *in vitro* culturing of mouse embryos using cavities of varying widths, has shown that both the geometry and the stiffness of the confining environment are important in driving and supporting fundamental processes such as axis polarization and cell lineage differentiation [28].

In this work, we confine regenerating Hydra in narrow cylindrical channels. Applying this external mechanical constraint during regeneration affects both tissue geometry and actomyosin fiber dynamics, enabling us to probe the interaction between them and their effect on the regeneration process and morphological outcome of the regenerated animal. We use patterned agarose gels to generate the confining channels, and introduce the samples into the channels using flow. We find that under anisotropic confinement, the regenerated body axis preferentially forms along the channel, in the direction of the ‘easy-axis’ of deformation imposed by the constraint. However, frustration between the inherited body axis alignment and the ‘easy axis’ along the channel, often results in an abnormal body plan and multiple body axes. We correlate these results with the actomyosin fiber organization in the confined geometry, and show that when the inherited axis is aligned with the channel, the array of fibers forms a minimal number of +1 defects at the two opposing tissue poles and normal regeneration proceeds. In contrast, in the frustrated orientation, the initially disordered fiber regions are spread out, and the confined tissue spheroid develops a nematic fiber array with excess positive and negative defects, and regenerates into animals with multiaxial morphologies. The influence of confinement on the fiber organization in the parallel and perpendicular configuration and the corresponding defect configurations that emerge, can be recapitulated by our recently-developed biophysical model of regenerating Hydra, which considers the coupled dynamics of the nematic actomyosin fiber organization and a biochemical morphogen [23].

## RESULTS

### Anisotropic confinement of regenerating Hydra in narrow cylindrical channels using patterned agarose gels

Confinement in an anisotropic, uniaxial geometry can be used to restrict and direct tissue dynamics along an externally-specified axis. Our previous work has shown that the body axis of Hydra regenerated from excised tissue segments is aligned with the inherited body axis from the parent animal [19, 21]. This opens the possibility to generate frustration between the externally-imposed axis and the direction of the inherited axis. To realize this, we set out to develop a methodology to anisotropically confine regenerating Hydra, which poses a number of challenges. First, since Hydra tissues do not grow appreciably in size during regeneration, so will not grow to fill the confining cavity, the full-sized tissue has to be introduced into the cavity without damaging it. Second, the confining method should support tissue viability and regeneration over several days, while being sufficiently restrictive to prevent the highly flexible and dynamic Hydra tissue from escaping or reorienting over time. Finally, the experimental configuration should allow for high-resolution live imaging over the entire regeneration process.

To achieve such anisotropic confinement, we introduce tissue spheroids into pre-formed narrow cylindrical channels using pressure-generated flow (Fig. 1; Movie 1). The channels are cast from stiff 2% agarose gel, while the channel cavities are filled with soft 0.5% low-melting agarose gel that is introduced together with the tissue samples. The stiff gel defines the walls of the confining cavity whose width is controlled by the diameter of the glass capillary used to cast the channel (Fig. S1; Methods). This width is chosen to be narrow enough to impose a significant constraint on the confined tissue, yet wide enough to prevent excessive shear that can tear the tissue during the insertion step. Specifically, we use channels between 120-180 μm wide, which is considerably smaller than the typical diameter of free regenerating tissue spheroids (200-400 μm). In this way, the channels provide an ‘easy-axis’ for tissue dynamics along the channel, while restricting movement in the perpendicular direction (Fig. 1B-D). The soft gel inserted with the sample does not impose significant constraints on the shape as it is easily deformed by the regenerating Hydra, yet it reduces movement of the tissue along the channel which facilitates high-resolution imaging.

**Figure 1:**
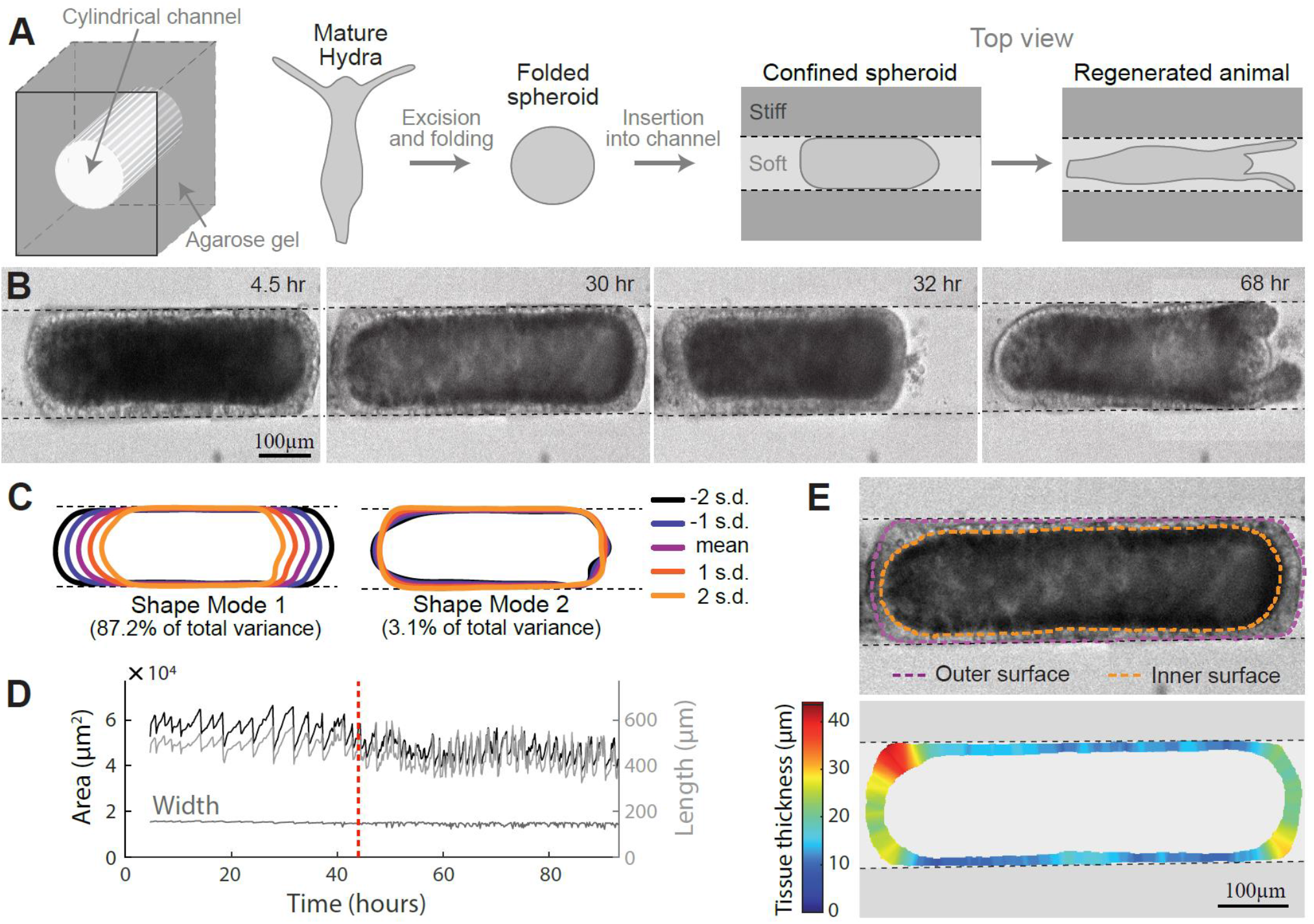
Hydra regeneration in narrow cylindrical channels. (A) Schematic illustration of the experimental setup. Left: 3D illustration of the channel geometry. Right: A tissue segment is excised from an adult Hydra and left to seal for 3-6 hours, and subsequently inserted into a cylindrical channel. The channel walls are made of stiff 2% agarose gel and the channel is filled with soft 0.5% low-melting agarose gel. The confined tissue spheroid regenerates within the channel. (B) Bright-field images from a time-lapse movie of a regenerating Hydra spheroid under confinement (Movie 1). The tissue geometry and shape dynamics are constrained by the channel, which limits the width of the confined spheroid as the tissue deforms, undergoes cycles of swelling and rapid deflations, and eventually regenerates. (C) Primary shape modes of a regenerating Hydra tissue extracted from the sample shown in (B). The average tissue shape is shown together with shapes one and two standard deviations away from the mean along each shape mode (Methods). The variation in tissue morphology is evidently restricted by the channel. (D) The projected area, length and width of the regenerating tissue as a function of time for the movie shown in B. While the projected area and the length are variable, the width is constrained and remains nearly constant. The regeneration time is 44 hours (dashed line). (E) Top: A bright-field image of a confined regenerating Hydra tissue spheroid. The inner and outer boundary of the bilayered epithelium are depicted. Bottom: the extracted tissue thickness determined from the normal distance between the inner and outer tissue boundaries.

Regenerating tissue samples are inserted into the channels using flow 3-6 hours after excision, allowing the excised tissues to complete their initial folding process and seal into hollow spheroids prior to their introduction into the channel (see Methods). The transient shear forces during the insertion step deform the tissues into elongated spheroids, but are typically sufficiently moderate to maintain the closed tissue morphology and prevent tearing of the tissue. The extent of deformation of the confined tissue in the channel depends on its size: inserted spheroids acquire an elongated ellipsoidal shape, filling the channel width, so that the tissue length along the channel, and hence its shape anisotropy, depend on the initial tissue size (Figs. 1, S2A-C). We find that despite the deformations and restrictions imposed by the channels, the vast majority of confined spheroids are able to regenerate in the channels (i.e., develop a visible mouth and tentacles), as in control conditions (65/78 in channels, compared with 141/166 for control tissues embedded in 0.5% low-melting agarose gel). Moreover, the regeneration times, defined as the time of tentacle appearance, are similar (42±13 and 37±11 hours, for confined and control samples, respectively).

Regenerating Hydra tissues are highly dynamic, exhibiting extensive, rapid shape changes throughout the regeneration process [19, 23]. These dynamics are primarily driven by contractions of the developing muscle fibers. In addition, Hydra spheroids undergo repeated cycles of gradual osmotic inflations and ruptures that lead to abrupt deflations [29, 30]. In the channels, we observe similar dynamics (Fig. 1D) but the overall morphological changes are essentially limited to changes in tissue length along the channel axis, whereas the tissue width is restricted by the channel diameter (Fig. 1). The regenerating tissue fills the width of the channel throughout the regeneration process, and only after the formation of a mouth and tentacles, does the body column become narrower than the channel width and separates from it. We also observe that the tissue sections that are pressed against the channel walls are thinner than those facing the open ends of the channel (and also thinner than in spheroids not confined in channels), likely due to the influence of the compressive counter-force exerted on the regenerating tissue by the channel walls (Fig. 1E).

To study the effect of confinement on Hydra morphogenesis, we follow tissue dynamics, actomyosin fiber organization, and regeneration outcome in confined Hydra tissues by live imaging. We use transgenic Hydra expressing Lifeact-GFP in the ectoderm and further introduce localized fluorescent cytosolic markers (sparse dextran labelling and/or photoactivated dye; see Methods) to track tissue movements [20]. The regenerating tissues are not adherent, so they are able to move relative to the channel walls. However, we find that tissue spheroids with an aspect ratio (length/width) larger than ∼2 largely maintain their orientation relative to the channel, whereas smaller tissues are able to rotate perpendicular to the channel axis (Fig. S4). As such, larger samples offer an opportunity to examine how the relative alignment between the inherited axis and the easy-axis of the channel impacts the regeneration process and in particular the axial patterning of the regenerating Hydra.

### Frustration between channel axis and inherited axis results in abnormal, multiaxial regenerated morphologies

The strong constraint on tissue shape and dynamics imposed by the channel is particularly interesting in the context of its effect on the body plan and outcome morphology of the regenerated Hydra. Our previous work showed that the inherited alignment of the supracellular actomyosin fibers guides the alignment of the body axis of regenerating Hydra [19, 20]. Excised Hydra tissues fold into spheroids containing a large domain of parallel fibers, aligned with the body axis of the parent animal, as well as disordered domains around the closure regions. Here we use the fiber alignment in the channel to obtain a readout of the inherited body axis in confined spheroids. In particular, this is useful for identifying frustration between the inherited body axis and the easy-axis of the channel when the primary initial fiber alignment is not aligned with the channel’s axis.

The relative angle between the inherited body axis and the channel axis depends on the orientation in which the tissue spheroid enters the channel. Since we use flow to drive the spheroids into the channel, the samples tend to orient with their long axis parallel to the flow direction, so the tissue spheroids typically become lodged along their long axis (Fig. 2A). The orientation of the inherited body axis relative to the long axis of the folded spheroid (prior to confinement) depends on the excised tissue geometry and the way it folds [19], and thus can be used to modulate the orientation of the inherited body axis relative to the channel axis upon confinement (Fig. S3). Specifically, spheroids originating from rectangular tissue segments tend to be oblong in relation to the fiber orientation, whereas spheroids originating from excised rings are typically oblate (Fig. S3A,B). Accordingly, once inserted into the channels, the majority of spheroids originating from rectangular tissues have their fibers primarily oriented along the channel direction, with a disordered domain that has a net topological charge of +2 [23], consisting of two cap regions at the far ends of the confined spheroid connected by a narrow bridge (Figs. 2B top, S3C). In contrast, spheroids originating from tissue rings are typically oriented with their fibers aligned mostly perpendicular to the channel, with two disordered domains, each with a net topological charge of +1 that are spread out along the walls of the channel, on opposite sides of the elongated tissue spheroids (Figs. 2B bottom, S3C).

**Figure 2:**
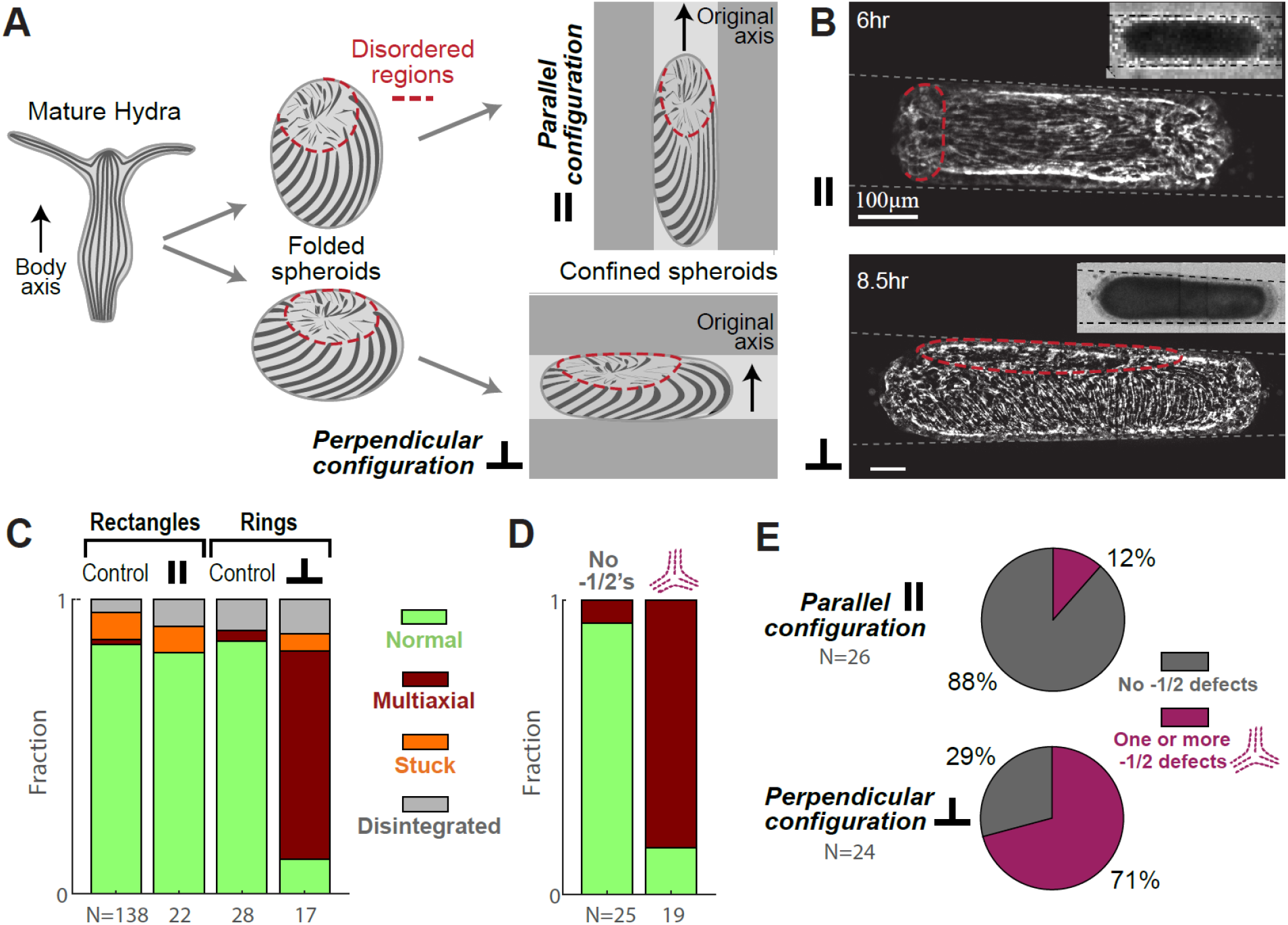
Initial orientation of tissue spheroids relative to the channel axis and its effect on morphological outcome. (A) Schematic illustration of the two main configurations of confined ellipsoidal tissue spheroids in channels. A spheroid originating from an excised rectangular strip will typically be oblong relative to its primary fiber orientation, and enter the channel in a parallel configuration. A spheroid originating from an excised ring will typically be oblate relative to its primary fiber orientation, and enter the channel in a perpendicular orientation. (B) Spinning disk confocal images of regenerating Hydra spheroids expressing Lifeact-GFP in the ectoderm following insertion into the channels in a parallel (top) or perpendicular (bottom) configuration relative to the channel. The visible disordered regions are indicated (dashed lines). (C) Bar plot showing the morphological outcomes of Hydra tissues with and without channel confinement. Tissues originating from rectangular strips confined in a parallel configuration regenerate normally with an average regeneration time of 38±15 hours (N=18 regenerated out of 22 samples, mean ± standard deviation), similar to those not confined in channels (37±10 hours; N=118 regenerated out of 138 samples). However, tissues originating from excised rings confined in a perpendicular configuration often regenerate into multiaxial morphologies with an average regeneration time of 44±12 hours (N=14 regenerated out of 17 samples, 12/14 multiaxial, 2/14 normal), unlike unconfined tissue rings which regenerate normally with an average regeneration time of 35±14 hours (N=22 with measured regeneration times, 25/28 samples regenerated). (D) Bar plot showing morphological outcomes of confined tissue spheroids as a function of whether -½ defects are observed (right; 16/19 multiaxial outcome) or not observed (left; 23/25 normal outcome) in the nematic organization of the actomyosin fibers during the first 24 hours following excision. The presence of -½ defects during the first 24 hours is correlated with higher prevalence of multiaxial morphology. (E) Pie plots showing the correlation between the initial alignment in the channels and the observation of -½ defects during the first 24 hours following excision.

We find that the inherited axis orientation and initial fiber alignment of the confined Hydra tissues relative to the channel’s axis have a striking impact on the regenerated body plan, in comparison to tissues of the same initial geometries without channel confinement (Fig. 2C). While tissues whose initial fiber orientation is aligned with the channel axis typically regenerate into a normal morphology, with their body axis aligned with the channel (i.e. one head and one foot along the channel axis, 18/18 of the regenerated animals Fig. S4A), samples that are initially oriented with their primary fiber alignment perpendicular to the channel direction often regenerate into multiaxial morphologies) 12/14 of the regenerated animals; Fig. S4B-D). Importantly, control Hydra spheroids that are embedded in 0.5% agarose gel without being confined in stiff channels regenerate with normal body morphology (Fig. 2C; >95% of regenerated animals), and this is true for the different excised tissue geometries used (rectangular segments or rings). The multiaxial morphologies arising from perpendicular-confined tissues consist mostly of animals with two heads (defined as a visible hypostome with a +1 defect in the nematic actomyosin fiber organization and at least one visible tentacle), and often with more than one foot or foot-like protrusions (Fig. S4B-D). Typically, these morphological features are not simply arranged along a single-axis, but rather include junctions between axes with characteristic -½ defects in fiber organization.

The effect of confinement on the regenerated morphology becomes more prominent with increased elongation of the tissue along the channel, which increases the extent of distortion generated by confinement (Fig. S2). Smaller tissues whose aspect ratios are closer to 1, are more likely to develop normal morphologies even when initially aligned perpendicular to the channel, in part because they are able to rotate and align along the channel. In contrast, the majority of larger tissues whose initial orientation is perpendicular to the channel, and are unable to rotate perpendicular to the channel’s axis, regenerate into abnormal morphologies (Fig. S2D).

### Formation of ‘excess’ defects under confinement is correlated with regeneration into multiaxial morphologies

How is the formation of multiaxial morphologies related to the initial orientation of the confined spheroids in the channel? To examine this, we follow actomyosin fiber dynamics from the initial configuration in the channel throughout the regeneration process. Over the first ∼24 hours of regeneration (both in normal conditions and under confinement), supracellular actomyosin fibers form in the domains that initially lacked ordered fibers, until the spheroid surface becomes fully covered with a parallel array of aligned nematic fibers interrupted only by point defects. Following this induction of order phase, the nematic field covering the surface must contain topological defects summing up to a total charge of +2 [20]. In the case of spheroids originating from rectangular tissue fragments and rings without channel confinement, the most common configuration of defects that arises is a +1 defect and a pair of +½ defects, that become the sites of head and foot formation, respectively [18, 20, 21].

Confined spheroids in the parallel configuration are found to form defects in the initially disordered caps at the far ends of the ellipsoidal tissue spheroid. Although these locations are hard to visualize because they are mostly curved away from the imaging plane, the fiber organization adjacent to these regions indicates that there is a net charge of +1 at each end (whether a single +1 or a pair of +½ defects), and no additional defects. We also observe localized tissue stretching and rupture events that are focused at the defect sites at the edges of the tissue (Fig. S5), exhibiting mechanical strain focusing during global fiber contraction events as seen in spheroids without channel confinement (see [23]). In this parallel configuration, we hardly observe -½ defects in the confined spheroids, and the tissues typically regenerate with a single axis and normal body morphology (Figs. 2C,E, 3). The parallel fiber orientation is maintained throughout the regeneration process, and the regenerated body axis forms along the ‘easy-axis’ of the channel (Fig. 3B,C).

**Figure 3:**
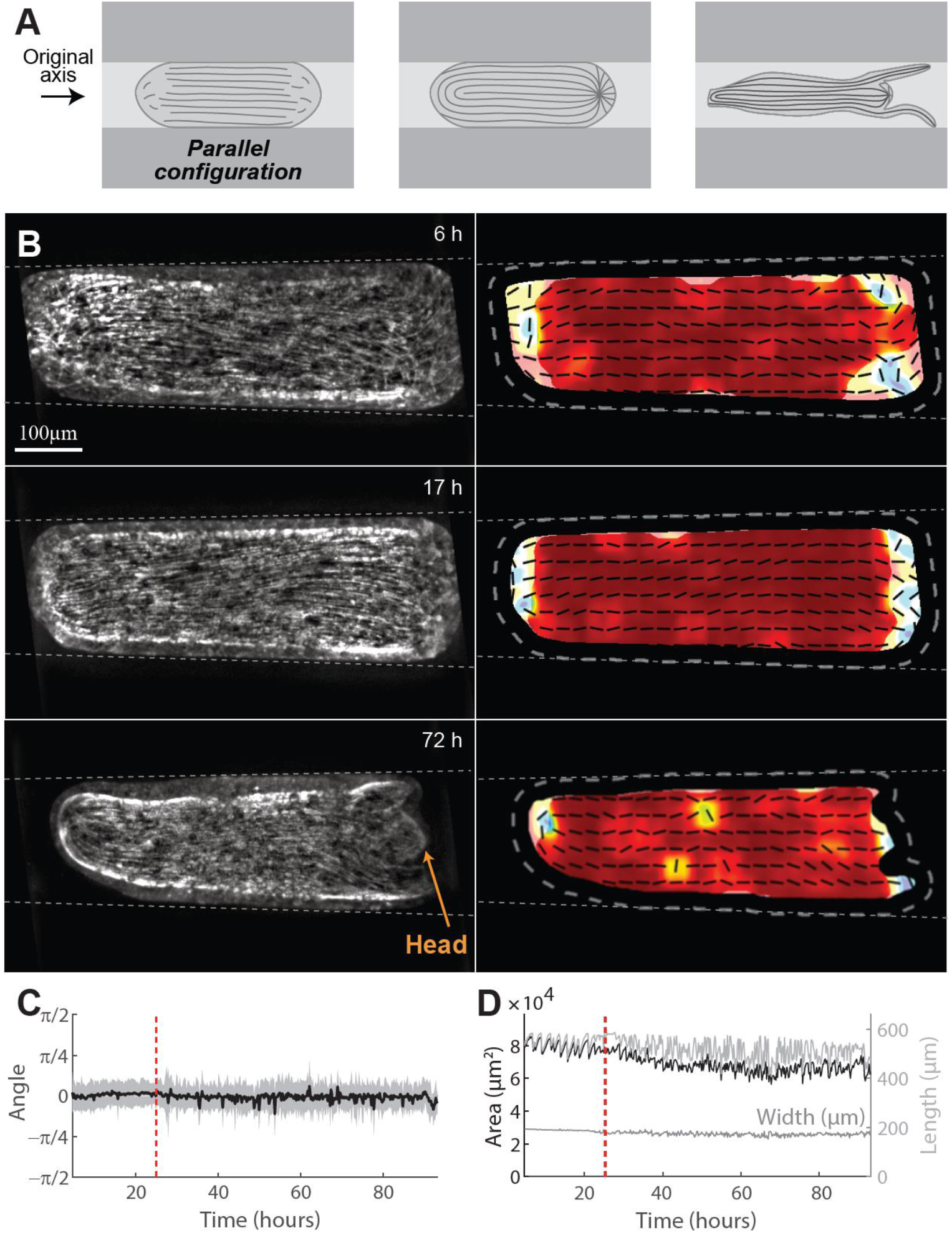
Dynamics of regeneration under channel confinement in the parallel configuration. (A) Schematic illustration of the fiber dynamics and defect configuration in confined spheroids in the parallel configuration. (B) Image series from a time-lapse movie of a regenerating tissue spheroid expressing Lifeact-GFP in the ectoderm (Movie 2). The fibers are initially mostly aligned parallel to the channel. The defects that localize at the sites of the future head and foot, emerge at the far ends of the tissue, and the spheroid regenerates with a single body axis aligned along the channel. Maximum projection images showing the actomyosin fiber organization (left), together with maps of the nematic orientation field (right; lines) and the local order parameter (color; see Methods). (C) Plot showing the observed fiber orientation (mean-black line, std-shaded region) relative to the channel axis for the regenerating sample shown in (B). The average fiber orientation is aligned parallel to the channel axis throughout the regeneration process. The regeneration time is 25 hours (dashed line). (D) Plots showing the projected tissue area, length, and width of the regenerating sample shown in B over time. The data shown in B-D are representative of N=18/18 rectangular tissues with initial parallel orientation that regenerated normally.

Contrary to this, in the case of frustrated spheroids, where the primary fiber orientation is perpendicular to the channel axis, the initially disordered domains are spread out, and we often observe the formation of multiple defects during the induction of order phase. Notably, this includes the presence of ‘excess’ positively-charged defects and an appropriate number of negative charges (-½ defects) to balance the extra positive charge (Fig. 2E). This defect configuration does not usually resolve (annihilation of positive and negative charges is rare), and the ‘excess’ positive charges typically become sites of additional heads or foot-like protrusions, whereas the -½ defects are found in the junctions between the emerging multiple body axes (Figs. 2D, 4, S4B-D). Here too, once +1 defects appear, they display repeated, localized swelling and rupture events, even when they reside along the channel walls, in the middle of the confined tissue (Fig. S6). Over time, the primary actomyosin fiber alignment in these multiaxial morphologies also converges to a mostly parallel orientation along the channel (Fig. 4B,C).

**Figure 4:**
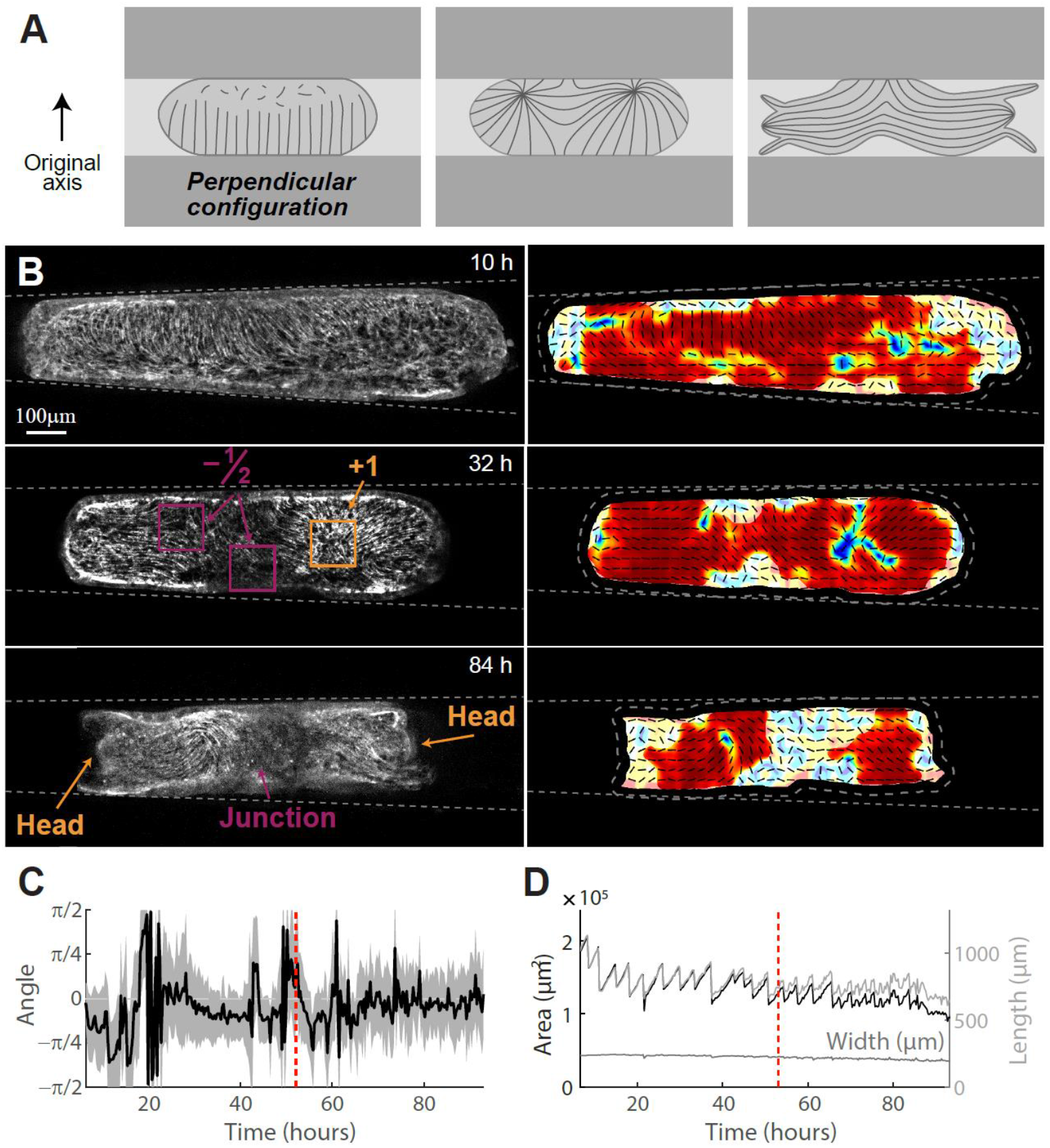
Dynamics of regeneration under channel confinement in the perpendicular configuration. (A) Schematic illustration of the fiber dynamics and defect configuration in confined spheroids in the perpendicular configuration. (B) Image series from a time-lapse movie of a regenerating tissue spheroid expressing Lifeact-GFP in the ectoderm (Movie 3). The initial configuration shows a region of fibers aligned perpendicular to the channel, and a spread out disordered domain (bottom right). Multiple defects, including -½ defects, emerge and the Hydra regenerates into an abnormal, multiaxial morphology (in this case, with two heads). Maximum projection images showing the actomyosin fiber organization (left), together with maps of the nematic orientation field (right; lines) and the local order parameter (color; see Methods). (C) Plot showing the observed fiber orientation (mean-black line, std-shaded region) relative to the channel axis for the regenerating sample shown in (B). The average fiber orientation is initially not aligned along the channel and the variability in local fiber orientations is high. Over time, the average fiber orientation becomes aligned with the channel, but the variability in local fiber orientations remains high. The regeneration time is 53 hours (dashed line). (D) Plots showing the projected tissue area, length, and width of the regenerating sample shown in B over time. The data shown in B-D are representative of N=12/14 ring tissues with initial perpendicular orientation that regenerated with multiaxial morphology.

### Biophysical model of Hydra regeneration recapitulates the effect of confinement on the emerging defect configuration

Overall, we find that confinement in cylindrical channels influences Hydra regeneration dynamics and morphological outcome, in a manner that is correlated with the changes in the defect configuration that emerge in the confined tissue spheroid (Figs. 2-4). When the inherited body axis and fiber alignment are parallel to the channel walls, the confined spheroids develop a defect configuration devoid of negatively-charged defects, and the Hydra regenerate along the channel’s axis with a normal body plan. However, when the early fiber alignment is primarily perpendicular to the channel, the configuration of defects that emerges typically contains multiple positive and negative defects. The excess defects typically persist and do not resolve, and are subsequently associated with the formation of multiaxial regenerated animals. To relate the defect configuration that emerges with the initial fiber pattern in the confined spheroids we turn to simulations of a biophysical model of Hydra regeneration dynamics (Fig. 5) [23].

**Figure 5.**
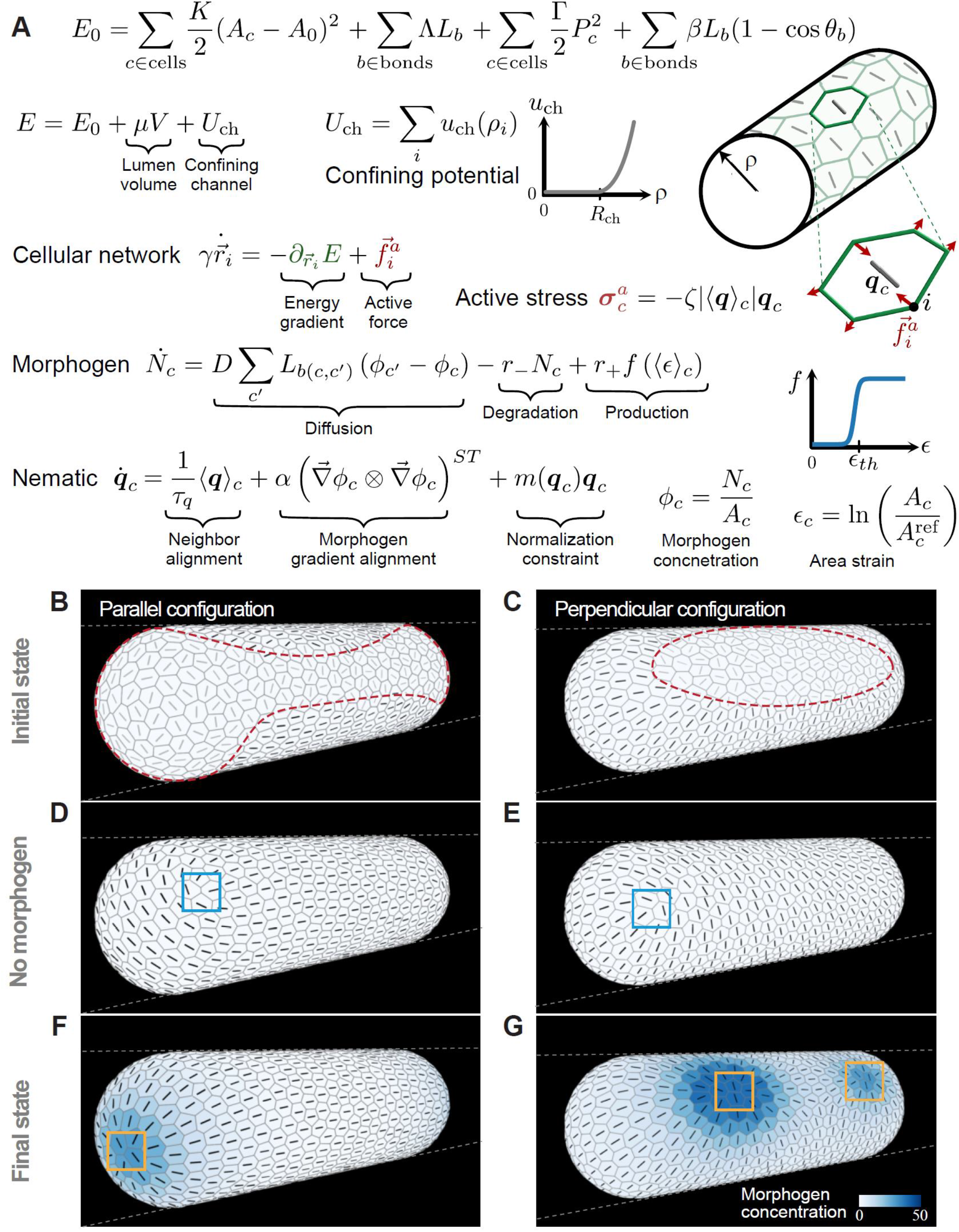
Biophysical model of Hydra regeneration in channels. (A) The model equations used to simulate the dynamics of Hydra regeneration in channels (Methods). The tissue mechanics are described by a vertex model energy on a deformable surface, E_0_, with added terms to reflect the constraints due to the incompressible lumen volume (V) and the channel confinement. A_c_ and A_0_ are the cell areas and preferred cell area, K is the area elastic modulus, Λ and L_b_are the bond tension and bond length, Γ and P_c_ are the perimeter elastic modulus and cell perimeter, β is the cell-cell bending modulus and θ_b_is the angle between normal vectors of neighboring cells that share the bond b. The confining potential u_ch_ penalizes cell vertices that move beyond the channel radius (R_ch_) from the channel axis. The total confining elastic energy is U_ch_ = ∑_i∈vertices_ u_ch_(ρ_i_), where ρ_i_ is the radial position of vertex i. An embedded nematic field (**q**_c_) and a morphogen concentration field (Φ_c_) are used to describe, respectively, the alignment of the contractile fibers and the morphogen concentration in each cell. The regenerating tissue dynamics are simulated using the dynamic equations for the vertex positions 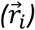, the nematic field (**q**_c_) and the number of morphogen molecules in a cell (N_c_). D is an effective transport coefficient that characterizes inter-cellular diffusion, L_b(c,c_′_)_ is the length of the bond shared by cell c and its neighbor c^′^, and ϕ_c_ = N_c_/A_c_ is the average morphogen concentration in a cell. r___ and r_+_ are the morphogen degradation and production rates, respectively. The morphogen production depends on cell area strain 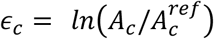, where 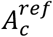 is the initial cell area at t=0. The production increases as a sigmoid function 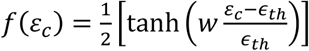, where ε_th_ is the strain threshold and w=10. The time-scale τ_q_ characterizes the nematic neighbor alignment rate, α characterizes the alignment of the nematic to morphogen gradients, and m(**q**_c_) is a Lagrange multiplier used to constrain the nematic magnitude to unity. For further details see Methods and Ref. [23]. (B,C) The initial nematic field organization used to initiate simulations of a confined tissue in the parallel (B) or perpendicular (C) configuration. The nematic in the initially disordered regions (dashed outline) is randomly orientated (gray). (D,E) Snapshots showing the nematic field organization in a confined tissue initiated in the parallel (D) or perpendicular (E; Movie 4) configuration in control simulations with no morphogen coupling after 30 contraction cycles. In both cases the tissue stabilizes in a four +½ defect configuration. (F,G) Snapshots showing the final nematic field organization and morphogen concentration after 30 contraction cycles in model simulations of a confined tissue initiated in the parallel (F) or perpendicular (G) configuration (Movies 5 and 6, respectively). The cell outlines, nematic orientations, morphogen concentrations and channel outlines are depicted in B-G.

We have recently shown that contraction of the supracellular ectodermal actomyosin fibers in regenerating Hydra, induces a characteristic pattern of mechanical strain focusing at defect sites [23]. A similar pattern of strain focusing at defect sites is seen in confined tissues, where the tissue at defect sites similarly experiences repeated, large stretching events during global fiber contractions (Figs. S5, S6). The observation of mechanical strain focusing and the emergence of morphological features at defect sites, led us to propose a biophysical model for Hydra regeneration that is based on a mechanochemical feedback loop, which couples the nematic field describing the actomyosin fiber alignment, the strain field in the tissue and a biochemical morphogen concentration field [23]. Our model proposes a self-organization mechanism whereby the mechanical strain focusing at defect sites is coupled to local production of morphogen. The fibers, in turn, align preferentially along the direction of the morphogen gradient, thus enhancing the strain focusing at defect sites, resulting in a positive-feedback loop that and promotes the establishment of a new head organizer (Fig. 5A). Here we use this model to simulate the influence of mechanical confinement on regenerating Hydra, and predict the different defect configurations that emerge in tissues confined in a parallel or perpendicular orientation relative the channel axis (Fig. 5B,C).

The model considers regenerating Hydra tissues as deformable sheets of cells enclosing a lumen, which are described by a vertex model forming a closed 2D surface (Fig. 5A) [23]. A nematic field describing the supracellular actomyosin fiber orientation and a scalar field describing the morphogen concentration are embedded on this cellular network. The time evolution of the model considers the dynamics of the nematic field, the tissue strain field and the morphogen concentration field as coupled fields. Muscle contractions during which the actomyosin fibers contract are included as pulses of active stress generated by the nematic field, motivated by our observations of large, recurring, global tissue contractions during regeneration [23]. The resulting tissue deformations induce strain-dependent morphogen production, and in turn, the nematic field aligns preferentially along morphogen gradients. Here we further extend this model to include the mechanical constraint due to confinement in the channel, by adding a confining potential to the energy function of the vertex model, mimicking the effect of the stiff channel walls on the tissue (Fig. 5A; see Methods for details).

We study the dynamics of the cellular network and the coupled nematic and morphogen fields within the confining potential. The embedded nematic field is initialized in two different configurations to reflect the initial actomyosin fiber organization in confined Hydra spheroids in the parallel or perpendicular configurations, respectively (Fig. 5B,C). Specifically, in the parallel configuration we consider a confined cellular surface with a large domain where the nematic field is aligned with the channel axis and a single disordered region that has a net topological charge of +2 (Fig. 5B). The disordered region consists of two disordered caps at the two far ends of the ellipsoid that are connected by a narrow disordered bridge. This configuration reflects the initial fiber organization in folded rectangular tissue pieces that become lodged with their inherited primary fiber orientation aligned with channel’s axis (Fig. 3). The perpendicular configuration is initialized as a cellular surface in which the fibers are aligned perpendicular to the channel axis, apart from two elongated disordered regions facing the channel walls, on opposite sides of the ellipsoid, each having a net charge of +1 (Fig. 5C). This fiber organization emulates the initial fiber pattern in confined spheroids generated from tissue rings, where the excised ring seals and forms two disordered regions on its top and bottom ends, that subsequently become stretched along the channel walls when the spheroid is introduced into the channel in a perpendicular orientation (Fig. 4).

Simulations of the model, which incorporates a mechanochemical feedback loop between the nematic organization, tissue strain and morphogen field, are able to reproduce qualitative features of the regeneration process under confinement (Fig. 5, Movies 5, 6), which are not reproduced if the feedback is not included (Movie 4). For initial conditions that correspond to the parallel configuration (Fig. 5B), we find that the actomyosin fiber organization develops two +1 defects at the centers of the two caps, coinciding with two local morphogen peaks that appear at these sites (Fig. 5F, Movie 5). The fibers in the initially disordered bridge become ordered along the channel axis, parallel to the fibers surrounding them. The tissue stabilizes in this configuration, with a parallel array of fibers aligned with channel’s axis and two +1 defects located at the far ends of the confined spheroid. This defect configuration agrees with our experimental observations in regenerating Hydra initiated in the parallel configuration that subsequently develop into a uniaxial Hydra with its polar body axis aligned with the channel (Fig. 3).

In contrast, in the perpendicular configuration, the actomyosin fiber organization develops excess defects (Fig. 5C,G, Movie 6). Each of the two initially disordered regions (with a net charge of +1 each) that are stretched along the channel walls (Fig. 5C), typically organize into two +1 defects near the shoulders of the ellipsoid, with additional -½ defects forming close to the central part of the disordered region (Fig. 5G). The resulting fiber configuration exhibits junctions with -½ defects near the center of the elongated spheroid, and multiple +1 defects facing the channel walls near the shoulders of the elongated ellipsoid. This defect configuration with excess +1 defects, together with additional -½ defects, emerging from the two initially disordered regions of the spheroid, mimics the experimentally-observed defect patterns in the perpendicular configuration (Fig. 4). Spheroids that develop this defect configuration typically regenerate into a multiaxial morphology, with additional heads and feet forming at the +1 defects sites located in the originally head and foot facing sides of the sealed ring, respectively (Fig. 4C). Note that simulations done in the absence of confinement predict a normal, uniaxial tissue morphology both for spheroids originating from folded fragments with a single disordered domain (with a net topological charge of +2) [23], and from sealed rings containing two disordered domains at either pole (having a net topological charge of +1 in each) (Fig. S7). As observed experimentally, the effect of confinement becomes more prominent as the aspect ratio of the confined tissues increases, with simulations predicting a transition from uniaxial to multiaxial configurations around an aspect ratio of 2 (Fig. S7). Notably, simulations that include the confinement but lack the coupling between the nematic and morphogen fields, predict a final configuration consisting of four +½ defects located at the “shoulders” of the ellipsoid for both the parallel and perpendicular initial configurations (Fig. 5D,E; Movie 4).

## DISSCUSSION

In this study, we demonstrate how confinement of regenerating Hydra tissues using patterned hydrogels can have a profound impact on their morphological outcome, and can lead to the formation of Hydra with multiple body axes under frustration. We relate this to the organization of the supracellular actomyosin fibers in confined Hydra, and the defect patterns that emerge in their nematic orientation. In the parallel configuration, two defect regions (with a total charge of +1 each) emerge at the two cap regions at the far ends of the elongated spheroid, and the animal regenerates with a uniaxial body plan aligned with the channel’s axis. However, in the perpendicular configuration, additional defects emerge along the length of the spheroid, with excess positive defects balanced by negative -½ defects. These frustrated spheroids, which exhibit excess topological defects following the induction of order in the fiber organization, develop into animals with multiaxial morphologies with the -½ defects localizing to the junctions between the different axes.

We suggest two main and interrelated effects that confinement can have on the regeneration process. The first effect is the strong influence on initial conditions. Insertion into narrow channels induces large deformations in the confined tissue, which in the perpendicular case, leads to substantial stretching of the domains lacking an ordered array of supracellular fibers along the channel walls. These domains are important because these are the regions in which the nematic defects in fiber organization form, and ultimately where the morphological features will emerge. The spread-out morphology of these disordered regions likely has an impact on subsequent fiber distribution and reorganization, and on the force generation within the tissue, as well as on the inherited polarity memory and biochemical signaling dynamics.

The second effect, following the initial deformation, is the influence of confinement within channels on tissue dynamics. The strong constraint on tissue geometry imposed by the channel walls must affect the patterns of mechanical stress experienced by the tissue, both due to the hydrostatic pressure gradient across the tissue and from the frequent muscle contractions that occur throughout the regeneration process [23]. Moreover, the stress pattern in the tissue during contractions will further be affected by the changes in fiber organization induced by the confining geometry. An indication that the dynamics within the channels affect fiber organization and regeneration dynamics is the strong preference of the body axes and the primary actomyosin fiber orientation to eventually align with the channel’s ‘easy-axis’. This alignment occurs even in the frustrated perpendicular configuration, whether by rotation if the tissue is sufficiently small or by formation of a convoluted body plan that appears as a ‘compromise’ between the inherited body axis and the preferred elongation axis and alignment along the channel (Fig. 4).

Our recently-developed biophysical model of Hydra regeneration provides a framework to analyze Hydra morphogenesis while considering the nematic actomyosin fiber alignment as a relevant field [23]. Even though the mechanistic details underlying the coupling between the morphogen field and the actomyosin fiber organization are still unknown, this framework enables us to recapitulate the qualitatively different defect configurations that emerge in confined spheroids initiated in the parallel and perpendicular configurations. Most notably, in the frustrated perpendicular configuration we show that excess defects, including negative defects, emerge (Fig. 5), as observed experimentally (Figs. 2,4). Spheroids that contain more defects typically develop into animals with a multiaxial body plan through subsequent morphogenetic events including tissue elongation and tentacle formation, which goes beyond the scope of our current model. The initial conditions used in the simulations mimic the initial nematic fiber organization observed in confined spheroids in the parallel and perpendicular configurations, respectively. However, we do not include the inherited asymmetry associated with the memory of polarity in the excised tissues, which has been shown to correlate with distinct morphogenetic events in the originally head and foot-facing sides of the tissues [21, 31]. Nevertheless, the model successfully relates the initial fiber organization with the formation of a single fiber axis, aligned with the channel, in the parallel configuration, and the emergence of multiple axes in the frustrated perpendicular configuration. As such, the model provides a useful framework for making predictions and examining the impact of confinement on the regeneration process.

The emergence of multiaxial morphologies appears to be a common hallmark of regeneration under frustration between inherited cues and tissue structure. This is apparent in regeneration from ring doublets [18], where frustration is created by fusion of two tissue rings with incompatible original polarities, in excised open rings which fold in a twisted manner [19], and here in the case of confinement in the perpendicular orientation. In all these cases, rather than one factor “winning” over the other, the frustration often results in a multiaxial morphology, indicating that both biochemical and structural cues, and the interplay between them, play a role in patterning the regenerated body plan.

Similar results have also been demonstrated recently by [32], who show that under compression of regenerating tissue segments, the deformation of the constrained tissue leads to a high prevalence of regenerated animals with multiaxial morphology.

The importance of early -½ defects as early indicators for the development of multiaxial morphologies is aligned with the previous observations of regeneration under frustration [18-20, 32]. While annihilation of -½ and +½ defects in the tissue spheroid is possible [20], more commonly the defect configurations that emerge in the folded spheroids following the induction of order remain relatively stable, and appear to predict the subsequent development of the body plan in the regenerating animal. In particular, the presence of early -½ defects is correlated with the appearance of multiaxial regenerated animal morphologies under various different perturbations [18-20, 32]. Interestingly, the specific pattern of excess defects that emerges varies depending on the characteristics of the initial conditions and constraints applied, and is correlated with the particular multiaxial body plan that develops. For example, the configuration with two heads on one side and two feet on the other that are connected by junctions at the middle, is common in confined tissues in the frustrated perpendicular configuration (Fig. 4) and consistent with the defect configuration predicted by the model (Fig. 5), yet is hardly ever observed in animals regenerated from excised open rings [19, 20] or frustrated ring doublets [18].

In summary, our results here show that mechanical constraints can lead to dramatic modulations of the body plan under frustrating conditions. The methodology developed for studying Hydra regeneration under uniaxial confinement can be further developed to introduce additional and possibly time-dependent constraints. Understanding the mechanisms through which confinement within narrow channels in the ‘frustrated’ orientation drives the formation of multiaxial morphologies remains an outstanding challenge. Nevertheless, the results presented here, together with additional recent results by us and others [18-21, 23, 30, 32], points towards the need for a more holistic approach for axial patterning in Hydra that integrates mechanics and biochemical signaling processes. Within this framework, local signaling events associated with axial patterning are dynamically coupled to the formation of topological defects in the actomyosin fiber organization through feedback loops, rather than being strictly “upstream” or “downstream” of each other [33]. More broadly, the appreciation of the importance of the synergistic interplay between biophysical and biochemical processes with extensive feedback across scales, implies that morphogenesis should not be considered a hierarchical, causal chain of events but rather a dynamic self-organization process.

## MATERIALS AND METHODS

### Hydra strains and culturing

The experiments are performed using transgenic Hydra strains expressing Lifeact-GFP in the ectoderm, generously provided by Prof. Bert Hobmayer (University of Innsbruck, Austria). Animals are cultured in standard Hydra culture medium (HM: 1 mM NaHCO_3_, 1 mM CaCl_2_, 0.1 mM MgCl_2_, 0.1 mM KCl, 1 mM Tris-HCl pH 7.7) at 18° C. The animals are fed 3 times a week with live Artemia Nauplii and rinsed after 4-8 hours.

### Regeneration experiments

Tissue segments are excised from the middle body section of mature Hydra, ∼24 hours after feeding, using a scalpel equipped with a #15 blade. Tissue rings are excised by performing two nearby transverse cuts. Rectangular tissue pieces are generated by first cutting tissue rings, and then cutting them further into three or four parts by additional longitudinal cuts.

Regeneration experiments are carried out at room temperature. Excised tissues are first allowed to fold into spheroids and seal in HM for 3-6 hours and are then embedded in 0.5% low gelling point agarose (Sigma) prepared in HM to reduce tissue movement during imaging. Control samples are made in 50mm glass-bottom petri dishes (Fluorodish). The samples are placed in few-mm sized wells that are cast of 2% agarose gel (Sigma) prepared with HM. The regenerating tissues are added to liquefied 0.5% low gelling agarose gel that is cool enough not to damage the samples (∼ 35°C), ∼ 3-6 hours after excision, and introduced into the precast wells and the soft 0.5% gel subsequently solidifies around the tissue. We add a few mm of HM from above ensure the gel does not dry out during imaging.

### Confinement in cylindrical channels

To study regeneration under anisotropic confinement, we embed regenerating Hydra in cylindrical channels made of agarose gel. The channel walls are made of 2% agarose gel, and the channels themselves are filled with low melting 0.5% agarose gel. The channels are prepared as follows. First, we use a manual puller to pull glass capillaries (1.65 mm outer diameter, 1.2 mm inner diameter; A-M systems) to create a tapered neck and long, narrow tip of approximately 120-180 μm width. The glass capillaries are placed in a custom-made Teflon mold with a narrow slit designed to hold the glass capillary in place, so that the tapering and narrow end of the capillary is suspended in a hollow cavity of around 0.5 cm in diameter (Fig. S1). The cavity is then filled with liquefied 2% agarose gel, which sets around the glass capillary. Once the gel has solidified, the capillary is removed by pulling backwards leaving a hollow cavity in the gel in the shape of the capillary. The same capillary is then truncated to remove the narrow end, and re-inserted into the gel, so the tapering end is aligned with the narrowest part of the channel in the gel and creates a funnel leading into the channel. A small hole is made from above using a narrower capillary at the far end of the channel, to prevent pressure build-up in the channel.

Hydra tissue segments are excised as described above and left to seal in HM for 3-6 hours. The tissues are added to liquefied 0.5% low gelling agarose gel that has cooled sufficiently, and inserted by flow through the glass capillary into the channel using a syringe. The inserted spheroids become lodged in the narrow end of the channel, and the soft gel that sets around them inside the channel prevents movement back into the wide section of the channel. The full agarose “block” containing the channel is then removed from the Teflon mold and mounted in a 50 mm glass-bottom petri dishes (Fluorodish) for imaging, with the channel aligned horizontally and as close as possible to the glass bottom for imaging.

### Tissue labelling

To label tissue regions we use either laser-induced activation of a caged dye (Abberior CAGE 552 NHS ester) or fluorescent dextrans that are electroporated into mature Hydra. Electroporation of the caged dye into live Hydra is performed using a homemade electroporation setup [20, 21]. The electroporation chamber consists of a small Teflon well, with 2 perpendicular Platinum electrodes, spaced 2.5 mm apart, on both sides of the well. A single Hydra is placed in the chamber in 10 μl of HM supplemented with 6-12 mM of the caged dye. A 75 Volts electric pulse is applied for 35 ms. The animal is then washed in HM and allowed to recover for several hours to 1 day prior to tissue excision. For laser-induced uncaging, the specific region of interest is activated following excision by a UV laser in a LSM 710 laser scanning confocal microscope (Zeiss), using a 20× air objective (NA=0.8). The samples are first briefly imaged with a 30 mW 488 nm multiline argon laser at up to 1% power to visualize the Lifeact-GFP signal and identify the region of interest for activation. Photoactivation of the caged dye is done using a 10 mW 405nm laser at 100%. The activation of the Abberior CAGE 552 is verified by imaging with a 10 mW 543 nm laser at 1%. Subsequent imaging of the Lifeact-GFP signal and the uncaged cytosolic label is done by spinning-disk confocal microscopy as described below.

Alternatively, we create a non-uniform pattern of marked cells in the body of a mature Hydra prior to excision, by electroporation of fluorescent dextrans. Electroporation of the probe into live Hydra is performed as described above, replacing the caged dye in the HM with a small drop of 0.2-0.3 μl of a cytosolic fluorescent marker (3kD dextran conjugated to Alexa fluor 647 or Texas Red at concentrations of 3 mM; Invitrogen) that is applied from above just before electroporation to generate an inhomogeneous dye distribution around the animal during electroporation, which leads to a non-uniform pattern of electroporated dye in the tissue.

### Microscopy

Spinning-disk confocal z-stacks are acquired on a spinning-disk confocal microscope (Intelligent Imaging Innovations) running Slidebook software. The Lifeact-GFP is excited using a 50 mW, 488 nm laser and the activated Abberior CAGE 552 is excited using a 50 mW, 561 nm laser. Images are acquired with an EM-CCD (QuantEM; Photometrix). Time-lapse movies of regenerating Hydra are taken using either a 10× air objective (NA=0.5), a 10x water objective (NA=0.45), or a 20x air objective (NA=0.75). Z-stacks are taken at 3μm intervals. The time resolution of the movies is 10-30 min. Imaging is done using appropriate emission filters at room temperature.

### Image processing and analysis

The tools used for image processing and analysis are based on a combination of custom-written code with adaptation and application of existing algorithms, written in MATLAB, Python and ImageJ, as detailed below [20, 23].

#### Image projections and identification of the tissue region

Analysis of the fiber orientation and tissue label is performed using 2D maximum projection images of the Lifeact-GFP and tissue label channels from 3D spinning disk stacks. Masks of the tissue are generated for every image based on the projected Lifeact-GFP signal, using automatic thresholding in ImageJ (‘Li’ method), and performing morphological operations (erosion, dilation, hole-filling) in MATLAB to obtain masks that accurately mark the tissue region in the image.

The thickness of the tissue within the channel is calculated by identifying the inner and outer surface outlines from the bright field images manually using the ‘magnetic lasso’ tool in Adobe Photoshop. A center-line is then determined from the inner and outer tissue contours, and the thickness for each point along the center-line calculated as the distance between the closest points to it on each outline.

#### Shape mode analysis

The projected shape outline of regenerating Hydra in the channels is extracted from the mask of the tissue and represented as polygonal outlines using the Celltool package developed by Zach Pincus [34]. The shape contours are described by a series of (x,y) points along the tissue boundary, which are resampled to 200 points that are evenly spaced along the boundary. The shape modes of the contours from a time-lapse movie are calculated by principal-component analysis using Celltool.

#### Fiber orientation and topological defect analysis

The local orientation of the supracellular ectodermal actomyosin fibers is described by a director field, which is oriented along the mean orientation determined in a small region surrounding every point in the 2D projected image. The nematic director field is determined from the signal intensity gradients in the projected images of the ectodermal fibers. Analysis of fiber orientation, number of defects, and relative alignment of fibers within the channel is determined manually, based on the calculated fiber orientation field and the local order parameter (as described in [20]) as well as the projected tissue label images that help with tracking tissue regions over time.

The initial alignment of the principle fiber orientation relative to the channel is classified into one of three categories - fully aligned with the channel, intermediate, and perpendicular to the channel. This is judged based on the visible supracellular fibers, and discounting the early ‘disordered’ regions where the nematic array of fibers is not yet formed. The judgement is performed ‘blind’ per sample, i.e. without knowledge of the final outcome.

The number of defects per sample are counted first by identifying defects in individual frames as described above. The samples rotate along the channel axis, so different frames in the movie can show the confined tissue from different orientations. We therefore use multiple images from the time-lapse movie to track defects between frames and identify the total number of visible defects. Since we cannot necessarily view the full surface of the sample, the number of defects identified provides a lower bound for the total number of defects present in the tissue.

#### Regeneration outcome analysis

Regeneration outcome in the channels is judged within a time window of 48-72 hours after excision. The regeneration outcome is classified into categories based on the number of feet/heads and any other protrusions observed, and how they are arranged relative to each other. Note that we do not functionally test for the presence of a foot, but judge the presence of a foot based on morphology. The different morphologies are then grouped into the following categories: normal (single axial morphology), multiaxial morphology, and those that failed to regenerate (separating between stuck - alive but with no observed tentacles or hypostome, and disintegrated).

### Model simulations

In our custom vertex model of Hydra tissue mechanics [23], individual cells are defined by the position of their vertices. Since the vertices belonging to a cell are in general not all in the same plane, we need to specify how we determine the cell area and cell tangent plane. To this end, we first define the cell center, 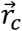, as the arithmetic average of all cell vertex positions. We then construct a collection of cell triangles by connecting every two neighboring vertices with the cell center. The cell area is defined as the sum of all cell triangle areas, and the normal vector to the cell tangent plane is defined as the area weighted average of the cell triangle normal vectors.

The vertex model energy function is defined as,

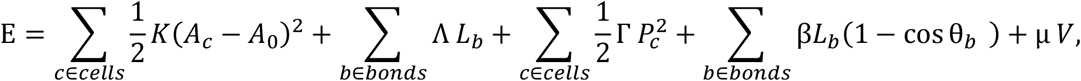

where *A*_*c*_ and *A*_*0*_ are the cell areas and preferred cell area, *K* is cell area elastic modulus, *Λ* and *L*_*b*_are the bond tension and bond length, *Γ* and *P*_*c*_ are the perimeter elastic modulus and cell perimeter, *β* is the cell-cell bending modulus and *θ*_*b*_is the angle between normal vectors of neighboring cells that share the bond *b*. The total volume of the lumen is denoted by *V*, and *μ* is a Lagrange multiplier that ensures lumen incompressibility.

To introduce the constraint associated with the channel, we introduce a cylindrical confining potential (Fig. 5A), such that motion of a cell vertex *i* beyond the channel radius *R*_*ch*_ costs an elastic energy *u*_*ch*_(*ρ*_*i*_) = (κ/2)(*ρ*_*i*_ −*R*_*ch*_)^2^Θ(*ρ*_*i*_ − *R*_*ch*_), where *ρ*_*i*_ is the radial distance between the vertex and the channel axis, κ is the channel elastic modulus, and Θ is the Heaviside theta function. We implement this potential as an additional contribution to the vertex model energy function,

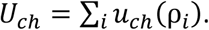

Each vertex position, 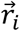, in the cellular network follows overdamped dynamics driven by the force associated with the gradient of the energy function 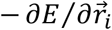 and an active force 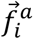 generated by the active stresses in all the cells to which this vertex belongs,

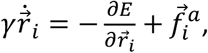

where *γ* is a friction coefficient. The active stress associated with each cell, 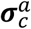, is generated by the contraction of the fibers in that cell upon activation. We introduce a unit nematic tensor ***q***_*c*_ in the tangent plane of each cell, representing the average orientation of the fibers in that cell. When cells move or deform, this nematic tensor is convected and co-rotated with them (see [23] for details). Based on the observations in Ref. [23] (see also Figs. S5,S6), we consider a sequence of global fiber contractions during the regeneration process. We emulate each global fiber contraction by simultaneously generating active stresses in all cells:

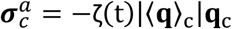

where the active stress magnitude *ζ*(*t*) is a Gaussian function of width Δ*T*_*ζ*_ and an amplitude *ζ*_*M*_. The contribution of each cell active stress to the active forces on its vertices is determined using the active stress scheme from [35], generalized to three dimensions.

The cellular network evolves according to the forces acting on the vertices. If at any time-step of the simulation, the length of any bond becomes smaller than a threshold value 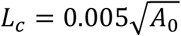, the two vertices of that bond are merged into a single vertex and the bond disappears. In this way 4-fold vertices are formed from merging of two 3-fold vertices. Such 4-fold vertices can resolve back into pairs of 3-fold vertices in one of two ways, reverting back to the original cellular configuration or rearranging the cells. A cell rearrangement event, also called a T1 transition, consists of the disappearance of one cell bond by the transient formation of a 4-fold vertex, that subsequently resolves into a new bond shared by cells that were previously not in contact. The stability of a 4-fold vertex is tested with respect to each of these two possible resolutions (see [23]). Note that our model also supports vertices with a larger number of bonds, but in practice we observe only 4-fold vertices in our simulations.

Following our observation that rearrangements are rare in the Hydra ectoderm, even during the large stretching events, we choose parameters of the vertex model so that the cellular network has a negative two-dimensional Poisson ratio (Table 1 and see Ref. [23]). This allows us to generate large isotropic strains without inducing a large number of cell rearrangements.

**Table 1:**
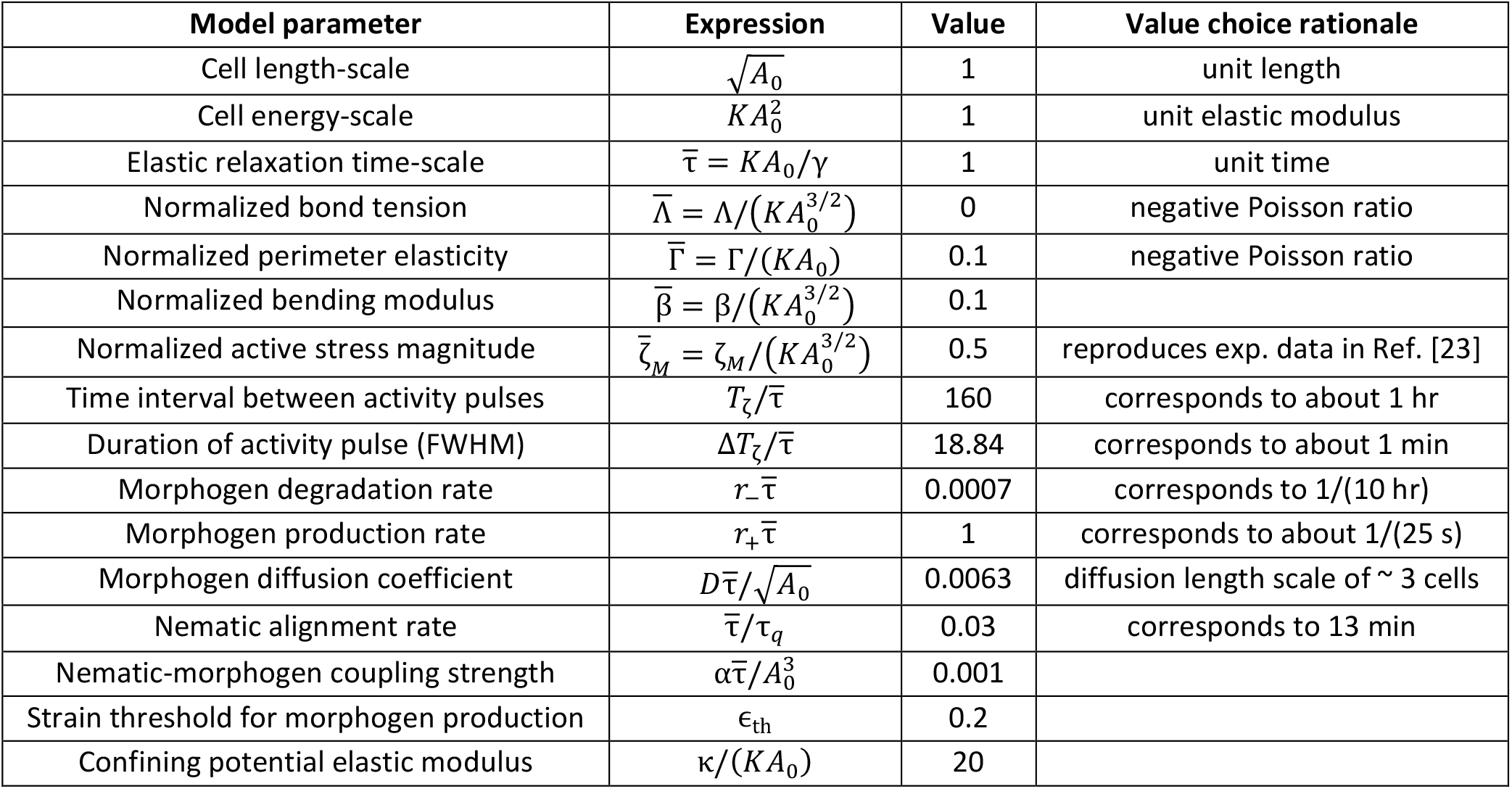
Values of the parameters used in the simulations. We summarize the rationale behind the choice in the last column, which is discussed in more detail in Ref. [23]. Correspondences to real times are based on experimental observations.

| Model parameter | Expression | Value | Value choice rationale |
| --- | --- | --- | --- |
| Cell length-scale | $\sqrt{A_0}$ | 1 | unit length |
| Cell energy-scale | $KA_0^2$ | 1 | unit elastic modulus |
| Elastic relaxation time-scale | $\bar{\tau} = KA_0/\gamma$ | 1 | unit time |
| Normalized bond tension | $\bar{\Lambda} = \Lambda/(KA_0^{3/2})$ | 0 | negative Poisson ratio |
| Normalized perimeter elasticity | $\bar{\Gamma} = \Gamma/(KA_0)$ | 0.1 | negative Poisson ratio |
| Normalized bending modulus | $\bar{\beta} = \beta/(KA_0^{3/2})$ | 0.1 | |
| Normalized active stress magnitude | $\bar{\zeta}_M = \zeta_M/(KA_0^{3/2})$ | 0.5 | reproduces exp. data in Ref. [23] |
| Time interval between activity pulses | $T_\zeta/\bar{\tau}$ | 160 | corresponds to about 1 hr |
| Duration of activity pulse (FWHM) | $\Delta T_\zeta/\bar{\tau}$ | 18.84 | corresponds to about 1 min |
| Morphogen degradation rate | $r_- \bar{\tau}$ | 0.0007 | corresponds to 1/(10 hr) |
| Morphogen production rate | $r_+ \bar{\tau}$ | 1 | corresponds to about 1/(25 s) |
| Morphogen diffusion coefficient | $D\bar{\tau}/\sqrt{A_0}$ | 0.0063 | diffusion length scale of $\sim 3$ cells |
| Nematic alignment rate | $\bar{\tau}/\tau_q$ | 0.03 | corresponds to 13 min |
| Nematic-morphogen coupling strength | $\alpha\bar{\tau}/A_0^3$ | 0.001 | |
| Strain threshold for morphogen production | $\epsilon_{th}$ | 0.2 | |
| Confining potential elastic modulus | $\kappa/(KA_0)$ | 20 | |

We introduce a morphogen field in our model as in Ref. [23]. The amount of morphogen in a cell *c* is described by the number of molecules *N*_*c*_ and develops over time according to,

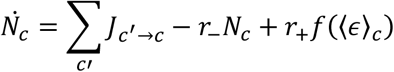

The first term reflects the contribution of diffusive fluxes between neighboring cells, that are of the form *J*_*c*_′_→*c*_ = *DL*_*b*(*c*,*c*_′_)_(*ϕ*_*c*_′ − *ϕ*_*c*_), where *D* is an effective diffusion coefficient that characterizes inter-cellular transport, *L*_*b*(*c*,*c*_′_)_ is the length of the bond shared by cell *c* and its neighbor *c*^′^, and *ϕ*_*c*_ = *N*_*c*_/*A*_*c*_ is the average morphogen concentration inside the cell. In this model intracellular diffusion is assumed to be much faster than transport between cells, so the morphogen concentration at the interface between two cells can be approximated by the average cellular concentration, *ϕ*_*c*_. We assume that the morphogen is degraded at a constant rate *r*_−_, and produced at a rate that is an increasing function of the cell area strain 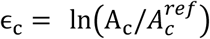, where 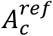 is the initial cell area at t=0. We use a sigmoid production function *f*(ϵ_*c*_) = 1/2 tanh(*w*(ϵ_*c*_ − ϵ_*th*_)/ϵ_*th*_), with *w* = 1*0*, so that *f* increases sharply as a function of the strain around the threshold strain value ϵ_*th*_. Therefore, the production is small below ϵ_*th*_, and quickly saturates to *r*_+_ above it. Note that, without loss of generality, we can set *r*_+_ = 1 which corresponds to expressing *N*_*c*_ in units of *r*_+_/*r*_−_ . The morphogen dynamics equation is therefore characterized by a degradation time-scale τ = 1/*r* and a length-scale 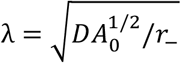 which reflects the length scale a morphogen diffuses before degrading.

To describe the dynamics of the nematic field, we assume that the nematic orientation in each cell is coupled to the nematic orientation in neighboring cells, and we further introduce a coupling of the nematic to the local morphogen concentration gradient,

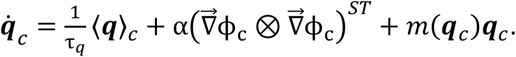

Here, the neighbor alignment is introduced through the average neighbor nematic operator ⟨***q***⟩_*c*_. On a flat surface, the average neighbor alignment would simply correspond to the arithmetic mean of the nematic tensors in neighboring cells. However, on a curved surface we need to account not only for in-plane alignment but also for the effect of surface curvature. In particular, since the nematic in each neighboring cell, ***q***_*c*′_, is constrained to the tangent plane of that cell, we first determine the projection ***P***_*c*_(***q***_*c*_′) to the tangent plane of cell *c* (see [23]), and then determine ⟨***q***⟩_*c*_ = ∑_*c*_′ ***P***_*c*_(***q***_*c*_′) /*n*_*c*_, where *n*_*c*_ is the number of neighboring cells. The dynamics of neighbor alignment is characterized by a time-scale τ_*q*_. To describe the alignment of the nematic field to the local morphogen gradient we first have to calculate the gradient on the curved surface. As mentioned above, we assume that the intracellular diffusion is fast compared to intercellular diffusion. To estimate the overall morphogen gradient across a cell we have to take into account the morphogen concentrations in neighboring cells. For each cell *c*, we aim to find a gradient vector 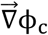 that satisfies the relation 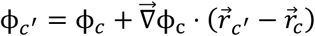 for all its neighboring cells *c*^′^. However, in general these relations cannot all be satisfied with a single vector and, so we define the morphogen gradient to be the vector 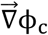 that produces the least-squared error of these relations (see [23]). Finally, due to the nematic symmetry, the nematic will align only to the axis of the morphogen gradient, independent of the polarity of the gradient along this axis. For this reason, the nematic aligns to a tensor constructed as the traceless-symmetric component of the outer product 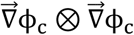. The parameter *α* characterizes the strength of alignment of the nematic to the morphogen gradient. Finally, we consider a nematic field with a fixed magnitude, describing the average orientation of the fibers in each cell. The nematic magnitude is maintained equal to 1 through a Lagrange multiplier *m*(***q***_*c*_).

We initialize the cellular network starting from a spherical tiling made of 800 hexagons and 12 pentagons, see Ref. [23]. We then introduce fluctuations of cell bond tension in each bond independently according to,

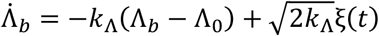

where ξ(*t*) is gaussian white noise: ⟨ξ(*t*)ξ(*t*^′^)⟩ = 2*D*_ξ_δ(t − t^′^), *k*_Λ_ = 1, *Λ*_*0*_ = −0.15, and *D*_ξ_ = 0.35. During the randomization process several cells become very small in area and are extruded so that we end up with 804 cells. After randomization we revert bond tension in all cells to the value used in simulations (see Table 1), and we minimize the vertex model energy to the nearest local minimum.

To constraint the cellular network in the channel potential we introduce a potential *u*_*ch*_(ρ_*i*_) = (κ/2)(ρ_*i*_ − *R*)^2^Θ(ρ_*i*_ − *R*) with *R* chosen to be slightly larger than the radius of the unconstrained network. Then we choose a particular value of the channel *R*_*ch*_ and we evolve 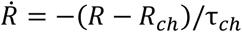 for 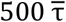, where we set 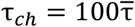, so that final *R* is within less of 1% from *R*_c*h*_. In the rest of the manuscript, we neglect this tiny difference and denote the channel radius by *R*_*ch*_. After constraining the cellular network by the channel potential, we relax any in-plane stresses introduced by the channel constraint by re-randomizing the cellular network following the same procedure described above.

The choice for all the model parameter values used in the simulations are given in Table 1, and the rationale behind these choices are presented in more detail in Ref. [23].

## ACKNOWLEDGEMENTS

We thank Erez Braun for many valuable discussions and comments on the manuscript. We thank Gidi Ben-Yoseph for excellent technical help. We thank Aurelien Roux and Yamini Ravichandran for sharing unpublished results and comments on the manuscript. We thank Alexandra Schauer, Jana Fuhrmann and Alex Mogilner for their comments on the manuscript. We thank Nitzan Dahan from the LS&E Imaging Center for advice and help in imaging.

This work was supported by a grant from the European Research Council (ERC-2018-COG grant 819174) to KK.

