## Supplementary Information for "Confinement modulates axial patterning in regenerating Hydra"

### Supplementary Figures

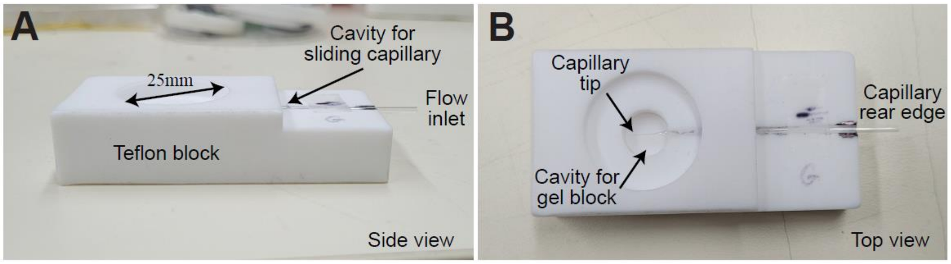


**Figure S1. Channel preparation setup.** Images of the Teflon block used for creating a cylindrical channel and loading the sample into it, from a side view (A) and a top view (B). A glass capillary that has been pulled to create an elongated tip with a diameter of 120-180 µm is inserted into a designated cavity in the Teflon block. 2% agarose gel is pipetted into the indicated region, so that it surrounds the capillary tip, and is then covered by a coverslip. After the gel hardens, the coverslip is removed, and the glass capillary is carefully pulled backward and its tip is broken. The capillary is loaded with the Hydra sample in liquefied 0.5% low melting agarose, reinserted into the cast gel channel, and connected at its rear end to a syringe. The capillary is then used to introduce the Hydra tissue spheroid (embedded in 0.5% gel) into the channel by applying pressure with the syringe. After loading, the capillary is removed and the solid gel block is extracted from the holder and placed on a coverslip for imaging.

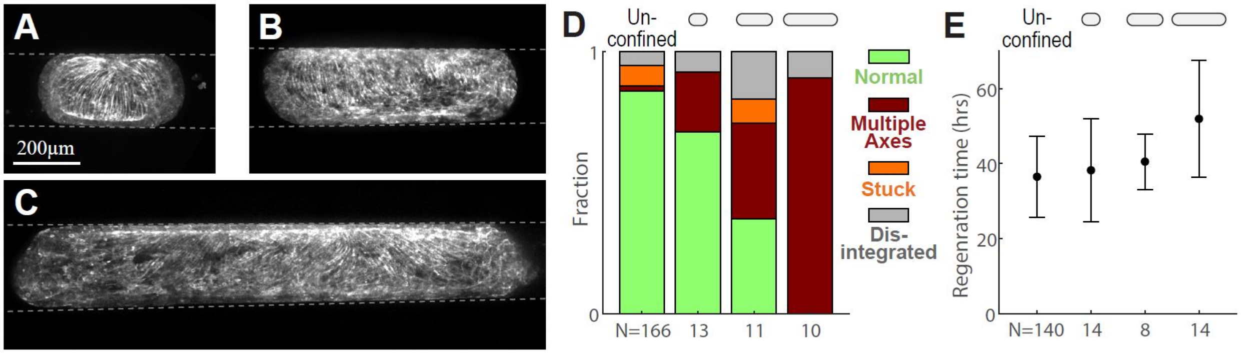


**Figure S2. The effect of perpendicular confinement on outcome morphology as a function of the aspect ratio of the confined tissue.** (A-C) Images of confined Hydra tissue spheroids in the perpendicular configuration, expressing Lifeact-GFP in the ectoderm. The confined tissues form cylindrical shapes with a small (A), intermediate (B) or large (C) aspect ratio. All images are maximum projections of spinning disk confocal stacks. (D) Bar plot showing the morphological outcomes of Hydra tissues without channel confinement and confined tissues in the perpendicular configuration with small (<2), intermediate (2-4) or large (>4) aspect ratios. Multiaxial morphologies are more prevalent for confined tissues that have a large aspect ratio. Note that confined tissues with a small aspect ratio often rotate and change their orientation relative to the channel. (E) The mean and standard deviation of the regeneration times (defined as the time of visible tentacle emergence) of Hydra tissues as in (D).

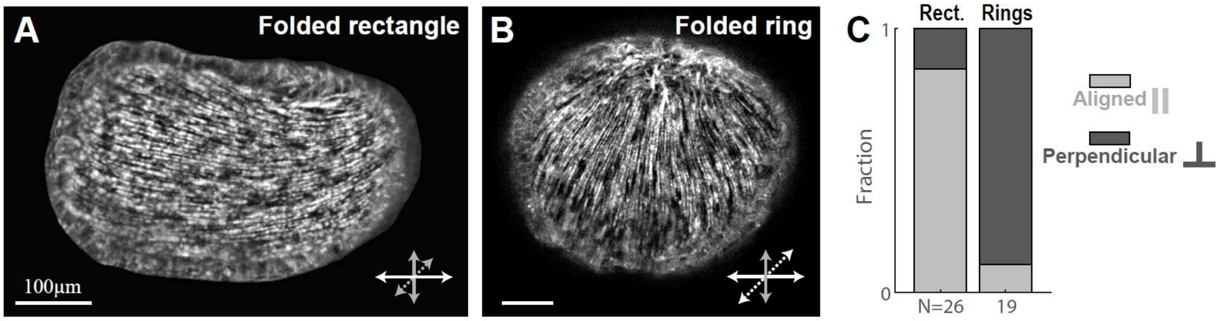


**Figure S3. Dependence of confinement configuration on excised tissue geometry.** (A,B) Images of Hydra tissue spheroids expressing Lifeact-GFP in the ectoderm in solution ~4 hours after excision. The spheroid originating from an excised rectangular strip is oblong relative to its primary fiber orientation (A), whereas the spheroid originating from an excised ring is oblate relative to its primary fiber orientation (B). (C) Bar plot showing the confinement configuration of spheroids originating from rectangular tissue pieces or tissue rings. Spheroids originating from excised rectangular pieces primarily enter the channel in a parallel configuration, whereas spheroids originating from excised tissue rings typically become lodged in a perpendicular orientation.

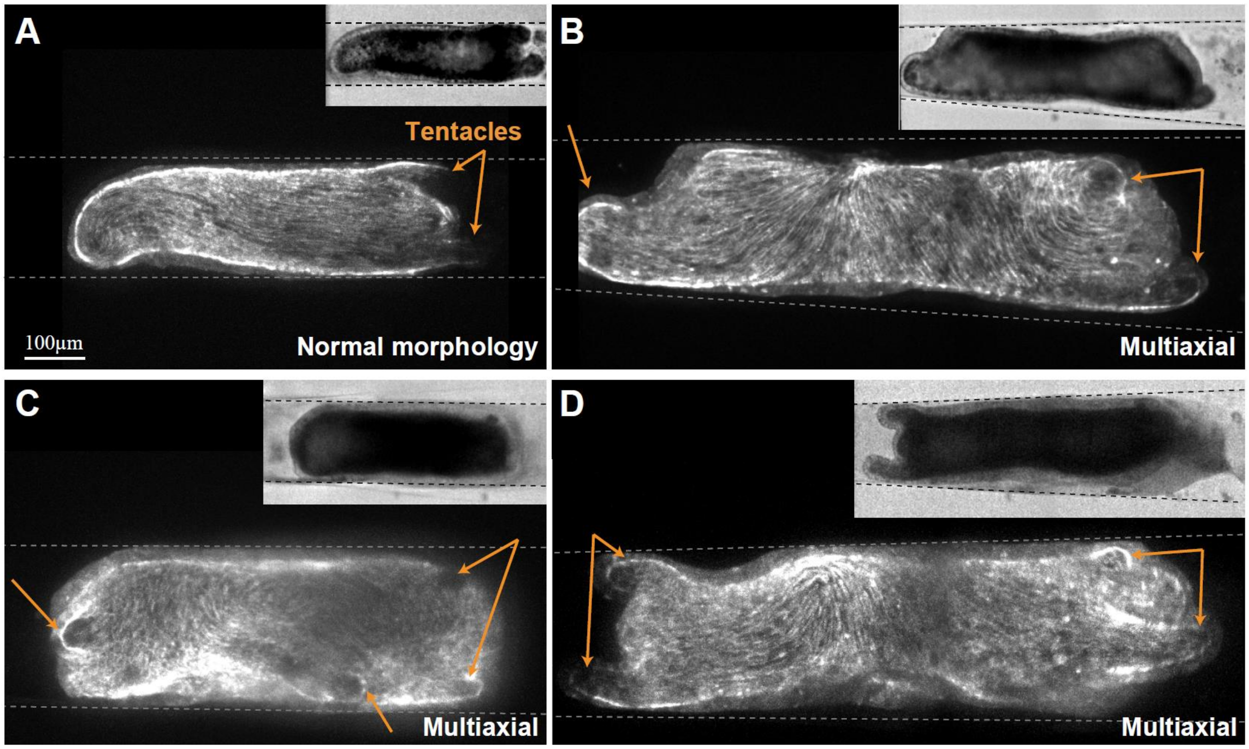


**Figure S4. Outcome morphologies in confined spheroids.** (A-D) Images of outcome morphologies of confined Hydra tissues expressing Lifeact-GFP in the ectoderm 48-96 hours after excision. The samples were confined in a parallel (A) or perpendicular configuration (B-D). The regenerated Hydra exhibit normal morphology (A) or multiaxial morphologies (B-D) with more than one region with tentacle formation (arrows). The images shown are maximum projections of spinning disk confocal stacks or bright-field images (inset)

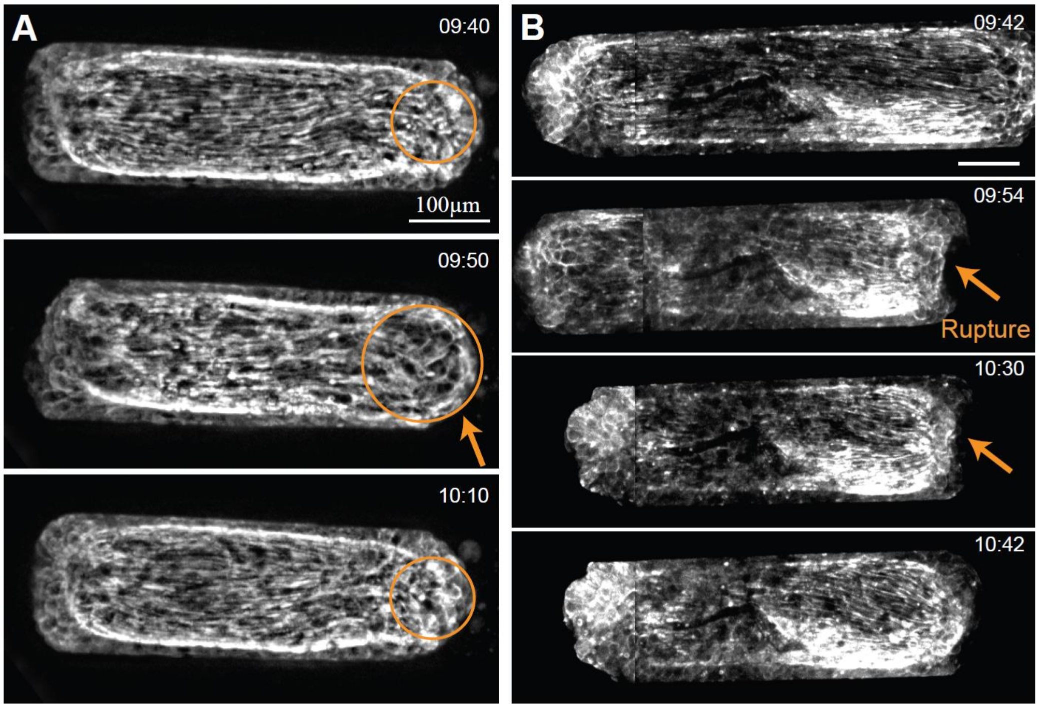


**Figure S5: Local tissue stretching and rupture events in confined spheroids in the parallel configuration**. Image series of confined regenerating Hydra spheroids in the parallel configuration expressing Lifeact-GFP in the ectoderm. Defect regions with a total charge of +1 are located at the two far ends of the tissues. Local tissue stretching (A) and rupture events (B) at defect site are indicated. The mechanical strain focusing induced by actomyosin fiber contraction in the defect regions in the confined tissue is similar to our observations in Hydra tissue spheroids without channel confinement. All images are maximum projections of spinning disk confocal stacks, and the elapsed time from excision is indicated (hh:mm).

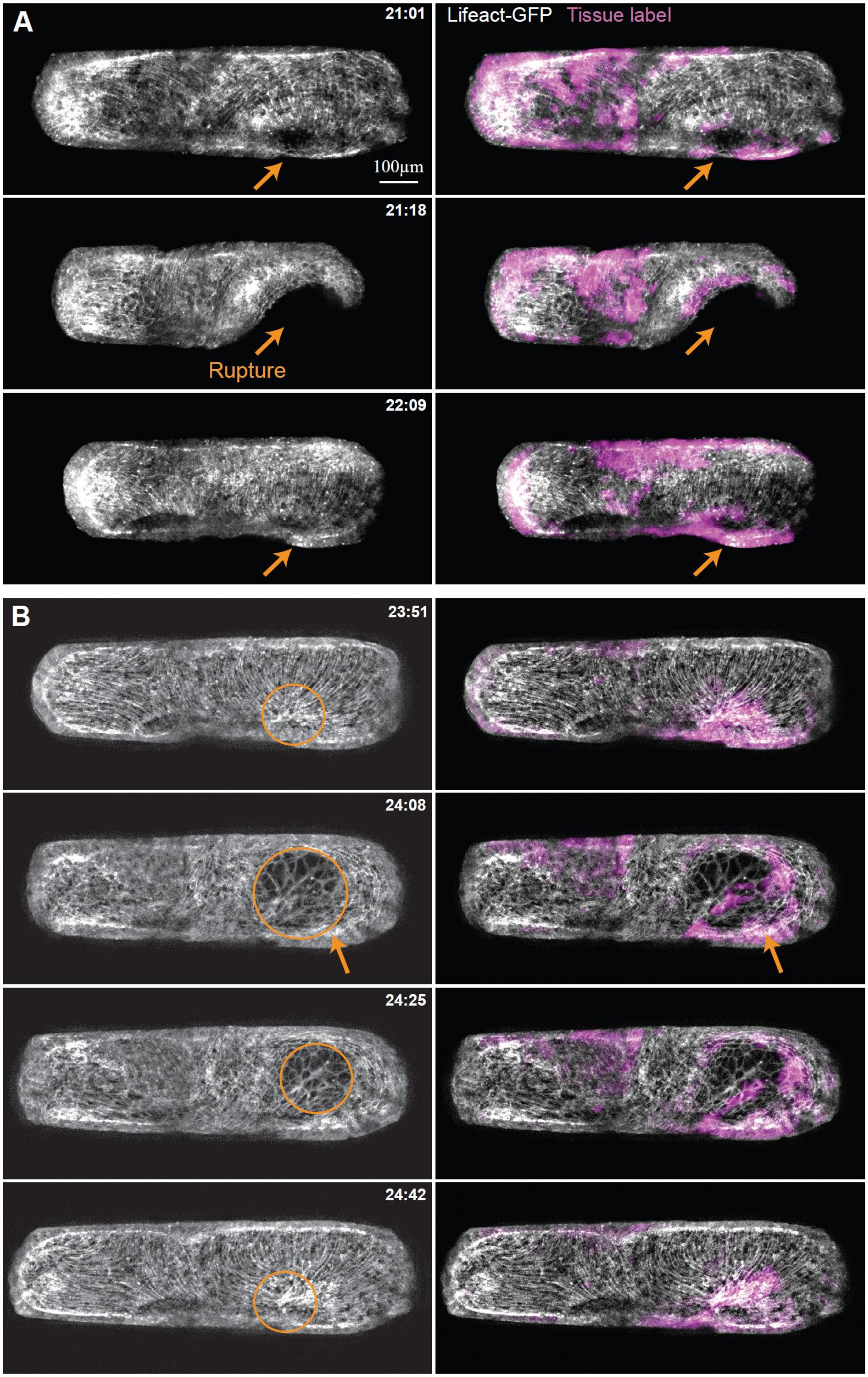
A
21:01
Lifeact-GFP
Tissue label
100μm
21:18
Rupture
22:09
B
23:51
24:08
24:25
24:42

**Figure S6: Local tissue stretching and rupture events in a confined spheroid in the perpendicular configuration**. Image series of a regenerating Hydra spheroid confined in a perpendicular configuration expressing Lifeact-GFP in the ectoderm (Movie 3). Multiple defects form, including in this case an excess +1 defect that is facing the channel wall. This +1 defect region undergoes local tissue stretching (A) and rupture (B), exhibiting mechanical strain focusing induced by fiber contraction as observed in Hydra tissue spheroids without channel confinement. Maximum projection images depicting the fiber organization (left), are shown together with an overlay of a laser-induced uncaged tissue label (magenta; Abberior CAGE 552, Methods). The fluorescent tissue label enables us to track the group of cells at the defect site during these large deformations. The elapsed time from excision is indicated (hh:mm).

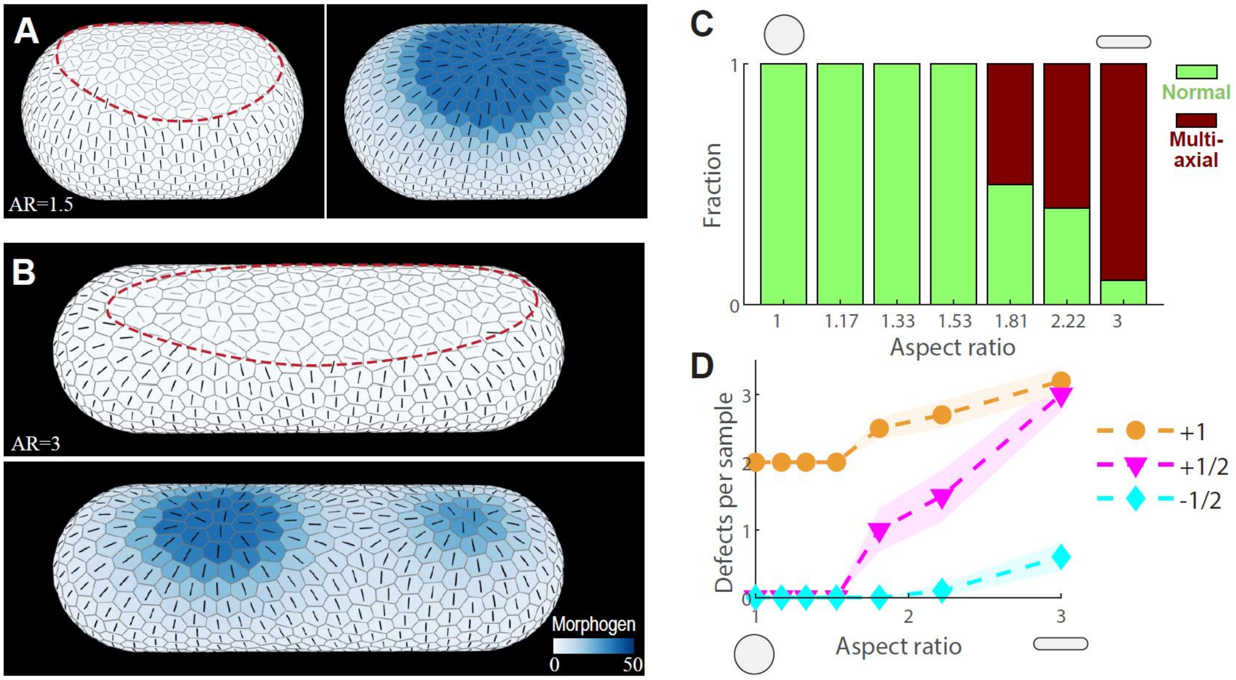


**Figure S7. Simulations of the effect of perpendicular confinement on fiber organization as a function of the aspect ratio of the confined tissue.** Regenerating Hydra tissues are simulated using the dynamical equations for the vertex positions, the nematic field and the morphogen concentration, with a cylindrical confining potential (Fig. 5; Methods). The aspect ratio of the confined tissue is varied from 1 to 3 by changing the diameter of the confining potential, while keeping all other model parameters fixed. The simulations are initiated in the perpendicular configuration with no morphogen. (A) The initial (left) and final (right; after 30 contraction cycles) nematic field organization and morphogen distribution of a confined tissue with an aspect ratio of 1.5 (Movie 7). (B) The initial (top) and final (bottom) nematic field organization of a confined tissue with an aspect ratio of 3 (Movie 6). The disordered region in the initial configurations is indicated (dashed line). (C) Bar plot showing the fiber organization in simulations of confined Hydra tissue spheroids initiated in the perpendicular configuration as a function of the aspect ratio of the confined tissue. (D) Graphs showing the average number of defects from each type, in simulations of confined Hydra tissue spheroids initiated in the perpendicular configuration as a function of the aspect ratio of the confined tissue (line: mean, shaded region: standard error of the mean). In C and D, 10 different realizations of the disordered region in the initial conditions are examined (see Methods), and the behavior of the simulations for the different realizations for each value of the confining potential diameter are shown.

### Supplementary Movie Captions

**Supplementary Movie 1: Hydra regeneration in narrow cylindrical channels**
Bright-field time-lapse movie of a regenerating Hydra spheroid under confinement (Fig. 1B). The tissue was excised and allowed to seal into a spheroid, before being inserted into a narrow cylindrical channel using flow (Methods). The geometry and shape dynamics of the regenerating tissue are constrained by the channel, which limits the width of the confined spheroid as the it deforms. The confined tissue spheroid undergoes cycles of swelling and rapid deflations and eventually regenerates into a mature Hydra, which in this example has a single body axis aligned with the channel. The elapsed time from excision is displayed (hh:mm), and the scale bar is 100 µm.

**Supplementary Movie 2: Dynamics of regeneration under channel confinement in the parallel configuration**
Time-lapse, spinning-disk confocal movie of a regenerating tissue expressing Lifeact-GFP in the ectoderm, introduced into the channel in a parallel configuration (Fig. 3). A rectangular tissue fragment was excised and allowed to seal into a spheroid, before being inserted into a narrow cylindrical channel using shear flow (Methods). Projected images show the supracellular ectodermal actomyosin fibers in the confined tissue (top) together with a map of the corresponding fiber orientation (bottom; black lines) and local order parameter (color shading; see Methods). The tissue outline (dashed line) and channel boundaries (solid lines) are indicated. The fibers are initially mostly aligned parallel to the channel. Defects emerge at the far ends of the confined spheroid, at the sites that will become the head and foot of the regenerated Hydra which forms a single body axis aligned with the channel's axis. The elapsed time from excision is displayed (hh:mm), and the scale bar is 100 µm.

**Supplementary Movie 3: Dynamics of regeneration under channel confinement in the perpendicular configuration**
Time-lapse, spinning-disk confocal movie of a regenerating tissue expressing Lifeact-GFP in the ectoderm, introduced into the channel in a perpendicular configuration (Fig. 4). A tissue ring was excised and allowed to seal into a spheroid, before being inserted into a narrow cylindrical channel using flow (Methods). Projected images show the supracellular ectodermal actomyosin fibers in the confined tissue (top) together with a map of the corresponding fiber orientation (bottom; black lines) and local order parameter (color shading; see Methods). The tissue outline (dashed line) and channel boundaries (solid lines) are indicated. The initial configuration shows fibers aligned perpendicular to the channel. Over time, multiple defects, including -½ defects (magenta), emerge within the initially disordered region, and the Hydra regenerates into an abnormal, multi-axial morphology, in this case, with two heads. The elapsed time from excision is displayed (hh:mm), and the scale bar is 100 µm.

**Supplementary Movie 4: Simulation of a Hydra tissue under confinement in the perpendicular configuration with no morphogen coupling.**

Simulation results showing a regenerating tissue spheroid confined in a narrow cylindrical channel in the perpendicular configuration. The simulation is initiated in a confined spheroid geometry with a partially ordered fiber configuration with fibers aligned perpendicular to the channel's axis, apart from two elongated disordered regions facing the channel walls, on opposite sides of the ellipsoid, each having a net charge of +1 (Fig. 5C). The tissue deformations and the dynamics of the nematic are modeled through 30 recurring global contraction activation events (the cycle is indicated) without including coupling to a morphogen field. The tissue self-organizes into a configuration with four +1/2 defects and without aster-shaped +1 defects (Fig. 5E).

**Supplementary Movie 5: Simulation of a regenerating Hydra tissue under confinement in the parallel configuration.**

Simulation results showing a regenerating tissue spheroid confined in a narrow cylindrical channel in the parallel configuration. The simulation is initiated in a confined spheroid geometry containing no morphogen and a partially ordered fiber configuration with a large domain where the nematic field is aligned with the channel's axis and a disordered region that has a net topological charge of +2 (Fig. 5B; Methods). The tissue deformations and the dynamics of the nematic and morphogen concentration fields are modeled within the confining potential through 30 recurring global contraction activation events (the cycle is indicated). The tissue self-organizes into a configuration with aster-shaped +1 defects colocalized with morphogen peaks at the two far ends of the tissue (Fig. 5F).

**Supplementary Movie 6: Simulation of a regenerating Hydra tissue under confinement in the perpendicular configuration.**

Simulation results showing a regenerating tissue spheroid confined in a narrow cylindrical channel in the perpendicular configuration. The simulation is initiated in a confined spheroid geometry containing no morphogen and a partially ordered fiber configuration with fibers aligned perpendicular to the channel's axis, apart from two elongated disordered regions facing the channel walls, on opposite sides of the ellipsoid, each having a net charge of +1 (Fig. 5C; Methods). The tissue deformations and the dynamics of the nematic and morphogen concentration fields are modeled through 30 recurring global contraction activation events (the cycle is indicated). Multiple defects, including four negative -½ defects (magenta) emerge within the initially disordered regions, and the tissue develops a multiaxial morphology with four aster-shaped +1 defects (yellow) that colocalize with morphogen peaks (Fig. 5G).

**Supplementary Movie 7: Simulation of a regenerating Hydra tissue with an aspect ratio of 1.5 under confinement in the perpendicular configuration.**

Simulation results showing a regenerating tissue spheroid confined in a perpendicular configuration within a wider cylindrical channel, so that the aspect ratio of the tissue is equal to 1.5 (Fig. S7A). The simulation is initiated in a confined spheroid geometry containing no morphogen and a partially ordered fiber configuration with fibers aligned perpendicular to the channel's axis, apart from two elongated disordered regions facing the channel walls, on opposite sides of the ellipsoid, each having a net charge of +1 (Fig. S7A left). The tissue deformations and the dynamics of the nematic and morphogen concentration fields are modeled through 30 recurring global contraction activation events (the cycle is indicated). The tissue develops two aster-shaped +1 defects that colocalize with morphogen peaks (Fig. S7A right).
